# EGFR and tyrosine kinase inhibitor interactions probed by hydrogen-deuterium exchange and mass spectrometry (HDX-MS)

**DOI:** 10.1101/2024.10.14.618277

**Authors:** Kumar D. Ashtekar, Mark A. Lemmon, Yuko Tsutsui

**Affiliations:** Department of Pharmacology, Yale University School of Medicine, New Haven, CT 06520, USA; Yale Cancer Biology Institute, Yale University West Campus, West Haven, CT 06516, USA; Yale Cancer Center, Yale University School of Medicine, New Haven, CT 06520, USA

## Abstract

EGFR is one of the primary drug targets for treating non-small cell lung cancer (NSCLC) patients carrying EGFR oncogenic mutations in the tyrosine kinase domain (TKD). Such patients typically receive tyrosine kinase inhibitors (TKIs) to inhibit aberrant activation of EGFR; however, together with the appearance of the TKI-resistant mutations, TKIs’ severe side effects often limit their clinical usage. To develop TKIs with the wild-type sparing effect, the wild-type structures bound to various TKIs ought to be characterized, though comparisons of such crystal structures do not show clear differences. To characterize subtle EGFR TKD structural changes upon TKI binding that cannot be gleaned from crystal structure comparisons, we employed HDX-MS. We show inhibitor-dependent EGFR dynamics that are displayed even among the TKD bound to chemically similar inhibitors. Such inhibitor-dependent structural changes appear to underlie TKI side effects and the selectivity of covalent inhibitors.

**Highlights:**

- EGFR shows TKI-dependent dynamics even if TKIs are structurally similar.
- The stability of the TKI-encounter complexes correlates with their side effects.
- Covalent TKIs disrupt the binding pocket of wild-type EGFR.
- The structure of the osimertinib-L858R/T790M complex is extremely rigid.

## Introduction

Epidermal growth factor receptor (EGFR) is a member of the receptor tyrosine kinase (RTK) family and is important for activating various biological signaling including MAPK, Ras/Raf, and PI3K/AKT.^1–3^ Oncogenic mutations in EGFR intracellular tyrosine kinase domain (TKD) result in aberrant receptor activation, leading to uncontrolled cell growth.^1–3^ Oncogenic EGFR TKD mutations are highly prevalent in non-small cell lung cancer (NSCLC) patients, with the frequency of the occurrence varying between 10-49 % of all NSCLC cases, depending on ethnic groups.^4,5^ Tyrosine kinase inhibitors (TKIs) are the first-line treatment for NSCLC patients and inhibit EGFR by occupying its ATP binding pocket located between the N- and C-lobes (Figures 1A and S1). The first-generation reversible TKIs include erlotinib, gefitinib, and lapatinib, were FDA-approved between 2004 and 2007 for treating NSCLC patients carrying EGFR oncogenic mutations while lapatinib was FDA-approved for treating metastatic breast cancer patients.^6–8^ It was soon discovered that within 12 months of the 1^st^ generation TKI therapy, cancer patients inevitably develop TKI resistance, with nearly 50 % of such cases due to an acquired T790M mutation in the hydrophobic pocket of the EGFR TKD.^5,9^ As a result, the second-generation covalent TKIs, afatinib, and dacomitinib, were developed; however, their side effects partly due to wild-type EGFR inhibition make them impractical to administer in clinical settings.^10,11^ Osimertinib is a third-generation covalent inhibitor and is the current front-line TKI therapy for patients carrying activating EGFR TKD mutations regardless of T790M mutation status.^12–15^ Both 2^nd^ and 3^rd^ generation covalent inhibitors form a covalent bond with C797 in the hinge region of the TKD (Figures 1A and S1) to prolong their residence time in the binding pocket; however, NSCLS patients eventually develop C797S mutation in the TKD,^9,16,17^ which makes these covalent inhibitors unreactive. To overcome C797S-resistant mutation, the development of fourth-generation inhibitors, which target allosteric sites of EGFR TKD, has been intensely pursued.^9,18–20^

**Figure 1:**
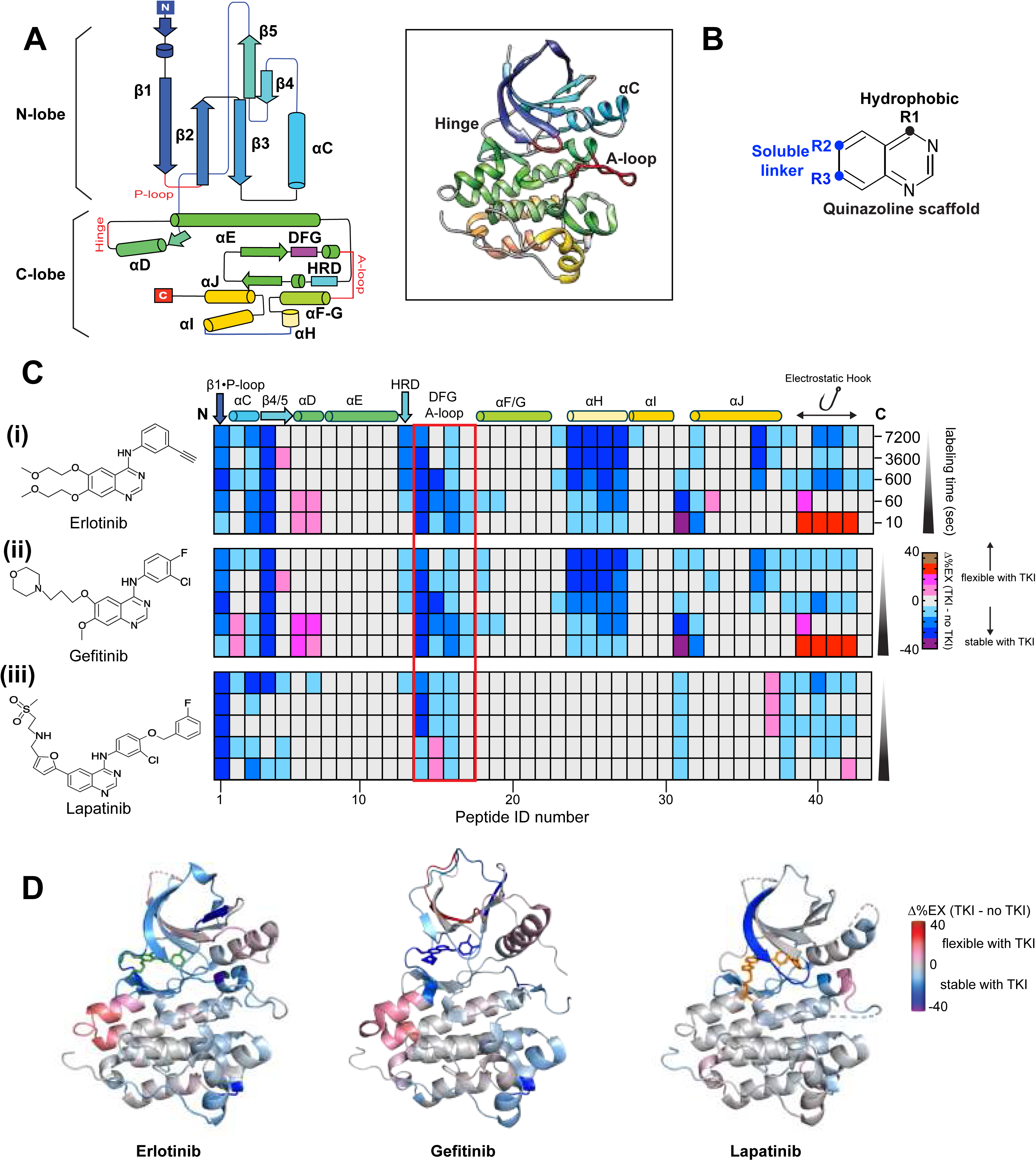
Structural dynamics of the wild-type EGFR TKD complexed with the first-generation TKIs probed by HDX-MS. **(A)** The 2D topology map of the EGFR TKD. The secondary structures and functionally important regions, including P-loop, hinge, HRD, DFG, and A-loop, are indicated. (inset) A crystal structure of the EGFR kinase domain (PDB ID: 4HJO)^25^ is shown. The secondary structures are colored in the same colors as the topology map. **(B)** The common quinazoline scaffold of the first-generation inhibitors with the position of the hydrophobic and soluble substituents indicated as black and blue-filled circles, respectively. **(C)** The Δ%EX heatmaps of erlotinib (i), gefitinib (ii), or lapatinib (iii)-bound wild-type TKD are shown. The columns show color-coded Δ%EX of common peptides in the inhibitor complexes at the indicated labeling times. Peptides are assigned peptide ID numbers from the N-terminus (left) to C-terminus (right) on the X-axis. *Supplemental information* lists the amino acid sequence and Δ%EX value of each peptide ID. The position of the secondary structures and conserved HRD and DFG motif regions are shown at the top of the heatmap. The red square box on the heatmaps indicates peptides corresponding to the DFG motif-containing binding pocket. **(D)** The Δ%Ex values of the indicated TKI-bound TKD at 1 min labeling time are color-coded as shown and mapped onto the EGFR TKD crystal structure bound to erlotinib (PDB ID: 4HJO),^25^ gefitinib (PDB ID:2ITY),^24^ or lapatinib (PDB ID: 1XKK).^6^ The bound TKI is shown as the stick in each figure. All HDX data were obtained from n = 3 separate protein preparations, with three independent experiments for each. HDX-MS data statistics are shown in Table S1. The structure maps are generated using PyMOL (http://www.pymol.org).

Numerous TKIs have been developed to discover a new class of TKIs with novel binding modes, and crystal structures of EGFR complexed with various TKIs have been solved.^21,22^ Based on these crystal structures, TKIs are mainly classified into two types, one selectively binding to the active EGFR structures with inward αC tilt and the other binding to the inactive EGFR with outward αC.^6,21,23–26^ MD simulation studies provide complementary structural information to crystal structures for studying the structural basis of TKI selectivity: Such studies typically assess how the free energy surface of EGFR TKD, which determines the population distribution of active and inactive structures, is reshaped by various oncogenic mutations.^27,28^ In addition, a quantum-mechanical simulation monitoring TKI binding processes at atomic details offers a possible structural basis for covalent TKI reactivity.^29^ Despite these intense computational efforts, structural mechanisms of TKI resistance have not been fully understood. As a result, a TKI with exquisite kinase selectivity without side effects has not been developed.

An ideal TKI producing desirable clinical outcomes should have wild-type sparing activity with no side effects. The discovery effort of such TKI typically involves positive design - the identification of inhibitors that maximize protein-TKI stabilities screened empirically using high-throughput thermal shift as well as kinase activity assays.^30,31^ Virtual inhibitor screening is also a common approach that identifies promising inhibitor hits by calculating various scoring functions using force-field-, empirical-, or knowledge-based methods.^32–34^ However, high-affinity binders do not always correlate with their specificity and selectivity.^35,36^ Rather, high affinity is often achieved at the expense of selectivity.^35,36^ Instead of searching for high-affinity inhibitors against a target kinase, the negative design could be an alternative approach for the TKI discovery effort. Negative design seeks to identify unwanted interactions and possibly destabilize such interactions to improve drug selectivity.^35,36^ In an effort to devise negative design-based TKI development, characterizations of unwanted interactions – interactions between wild-type kinases and various inhibitors are critical.

In this study, we probed interactions between wild-type EGFR TKD and various FDA-approved inhibitors by hydrogen-deuterium exchange and mass spectrometry (HDX-MS). We show inhibitor-dependent EGFR dynamics that cannot be categorized into the simple two-state inhibitor classification proposed by crystal structures of TKI-kinase comparisons.^21^ Our HDX-MS results also show a strong stabilizing effect of the second-generation TKIs on wild-type EGFR TKD encounter complex that may explain the structural basis of severe side effects. Finally, our HDX-MS results offer a possible structural basis for osimertinib clinical success against L858R and L858R/T790M mutants with the wild-type sparing effect.

## Results and Discussion

### Inhibitor-dependent dynamics are seen in distant regions from the binding pocket

The first-generation reversible TKIs, including erlotinib, gefitinib, and lapatinib, have the same quinazoline scaffold with variable hydrophobic moiety at R1 and soluble chemical groups at R2 and R3 substituents (Figure 1B). Comparisons of EGFR TKD crystal structures bound to these inhibitors show little structural differences with less than 1 Å backbone RMSD,^6,23–25^ despite differences in IC50.^24,37,38^ Such differences could be attributed to TKI-dependent changes in protein dynamics that are not seen in crystal structure comparisons.

To probe subtle structural changes upon the first-generation inhibitor binding to the EGFR TKD, we employed hydrogen-deuterium exchange and mass spectrometry (HDX-MS). EGFR TKD was pre-incubated with each inhibitor, and the inhibitor-bound TKD was labeled with a D_2_O-containing buffer for different time points. Structurally flexible or solvent-exposed amide hydrogens of the protein backbone readily exchange with deuterium at earlier labeling time points than amide hydrogens in structurally stable or buried regions.^39^ By comparing the deuterium uptake of the TKI-bound and -unbound TKD, regions that undergo structural changes upon the inhibitor binding are identified. Such comparisons at all deuterium labeling timepoints are expressed as Δ%Ex, with positive or negative Δ%Ex indicating regions that become structurally flexible or stable, respectively, upon the inhibitor binding relative to the TKD without inhibitors (Figures 1C, 1D, S2, and S3). Figure 1C shows comparisons of Δ%Ex of the same peptides derived from different TKI-bound TKD. The Δ%Ex values of all analyzed peptic peptides in each TKI-bound TKD are plotted as Woods plots in Figure S2. To aid visualization of inhibitor-dependent Δ%Ex in different structural regions, the Δ%Ex results are color-coded and mapped onto a crystal structure of the TKD (Figures 1D and S3). The HDX data statistics are presented in Table S1. The overall Δ%Ex pattern of the erlotinib- or gefitinib-bound TKD is highly similar in most of the secondary structure regions: Structural stabilization is seen in β1×P-loop, β4, the binding pocket containing the conserved HRD and DFG motifs (Figures 1C, 1D, S2-S4). Although the binding of both inhibitors destabilizes αD at the early labeling time points of 10 seconds and 1 min, a region N-terminal to αD, the hinge, shows strong stabilization at all labeling time points of seconds to hours (Figure S4). Erlotinib- or gefitinib-specific dynamics are seen in the N-terminal end of αC (residue 753-760), where structural flexibility of the αC region increases at 10 sec and 1 min only with gefitinib but not with erlotinib (Figures 1C, 1D, and S4). These results demonstrate that the dynamic responses to inhibitor binding involve timescale-specific structural changes that are unique to each inhibitor. In other words, TKI binding modulates the protein’s accessibility to exchange competent conformations in inhibitor-dependent manners.

The effect of erlotinib or gefitinib binding is propagated to the C-lobe, rigidifying the entire αH at all labeling time points (Figures 1C, 1D, S2-S4). The other C-lobe helices, including αF, the N-terminal region of αG, αI, and αJ also show structural rigidification, though in labeling time-dependent manners (Figures 1C, 1D, S2, and S3). A part of the C-tail containing D1006, D1009, and D1012 shows enhanced flexibility at 10 seconds labeling time point but becomes stabilized at later time points (Figure 1C). These acidic residues are thought to be important for forming a symmetric inactive kinase dimer via “the electrostatic hook,” salt bridges between the C-tail acidic residues and basic residues near the HRD and DFG motifs, including K823, K846, H850, and K852 that are seen in inactive TKD crystal structures.^40^ Although the C-tail Δ%Ex pattern containing the electrostatic hook is not direct evidence for the formation of the symmetric kinase dimer, the magnitude of the protection on 10-minute to hour timescales (Figure 1C) indicates this region to be involved in forming some ordered structural elements in the presence of the TKIs, consistent with inactive TKD crystal structures.^6,41^

Lapatinib is a dual inhibitor targeting both EGFR and HER2. The overall dynamics (Δ%Ex) of lapatinib-bound TKD show different patterns from that of the erlotinib- or gefitinib-bound TKD (Figures 1C, 1D, Figures S2-S4): The effect of lapatinib binding is confined mainly in the N-lobe, as the dynamics of the C-lobe helices are not impacted as much as erlotinib or gefitinib binding. The most notable differences are the dynamics of the αD and the HRD motif regions, which show no discernible structural changes except for the 2-hour labeling time point (Figures 1C, Figures S2-S4). Also, unlike erlotinib and gefitinib binding, destabilization of the A-loop region is evident at 10 sec and 1 min labeling time (Figures 1C, 1D, S2-S4). These results again demonstrate inhibitor-dependent dynamics.

Being an inactive state-selective inhibitor, lapatinib is expected to preferentially bind and stabilize the EGFR TKD with the αC-out conformation while gefitinib stabilizes the active αC-in conformation. Since the wild-type TKD predominantly populates the inactive αC-out conformation,^27,28^ the αC dynamics between the unliganded and the lapatinib-bound TKD are expected to be identical. However, time-dependent structural changes in αC and other regions (Figures 1C,1D, S2-S4) demonstrate that the TKD complexed with various inhibitors retains a propensity to deviate from ensemble-averaged crystal structures. This is important as protein dynamics drives allostery.^42^ Thus, our HDX-MS results show that structural states of a single or a few local regions cannot be a sole indicator for the functional state, in agreement with a general view of allostery – no single state necessarily dominates the ensemble because of proteins’ marginal stabilities and coupling energies.^43^

### The binding of chemically similar TKIs results in distinct EGFR TKD dynamics

The second-generation inhibitors are covalent inhibitors that form a covalent bond with Cys797 in the hinge region (Figures 1A and S1) via their acrylamide warhead. They include afatinib and dacomitinib, which have the same 3-chloro-4-fluoro aniline substituent at the hydrophobic end (R1) as gefitinib but carry different substituents at the aqueous-facing R2 and R3 positions (Figure 2A). These covalent inhibitors engage with the binding pocket reversibly before the covalent bond formation.^44,45^ If so, studying the dynamics of such encounter complexes is of prime interest for designing new covalent inhibitors that react efficiently and selectively with a target kinase. To gain insight into the structural dynamics of such encounter complexes by HDX-MS, we utilized non-covalent counterparts of these inhibitors that have the acrylamide group replaced with the amide moiety (Figure 2A and Figures S5-S10), which reversibly binds to the TKD but does not react with Cys797.

**Figure 2:**
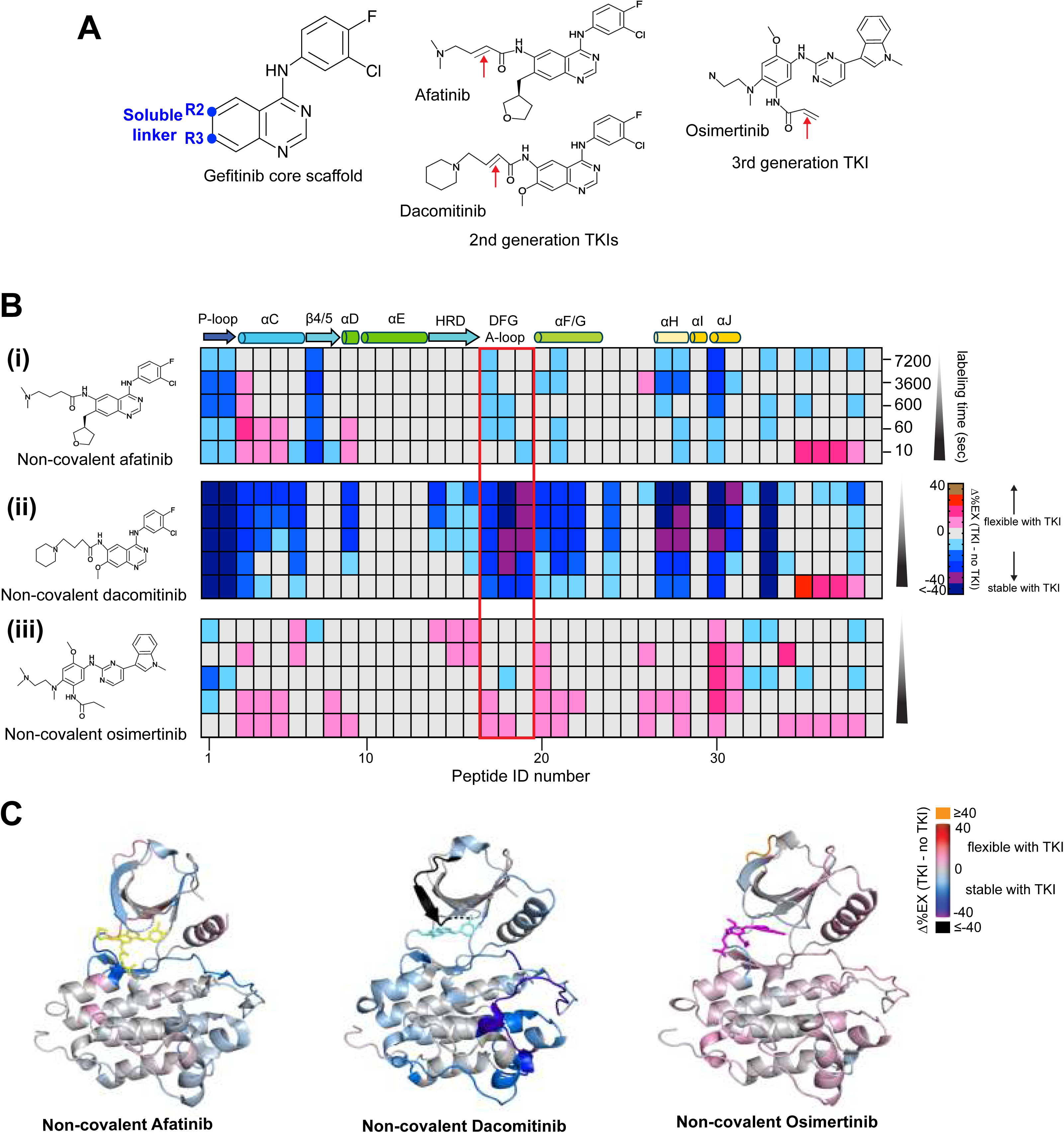
Structural dynamics of the wild-type TKD complexed with the non-covalent second- and third-generation TKIs probed by HDX-MS. **(A)** The second-generation TKI shares the common gefitinib core scaffold with variable soluble linker substituents at R2 and R3 positions, indicated as blue-filled circles (left). The chemical structure of the second- and third-generation inhibitors is shown with the red arrow indicating the position of a reactive acrylamide warhead. The warhead was replaced with the non-reactive amide as described in the *Methods* to probe the dynamics of wild-type EGFR with the reversible second- and third-generation inhibitors. **(B)** The Δ%EX heatmaps of non-covalent afatinib (i), non-covalent dacomitinib (ii), or non-covalent osimertinib (iii)-bound wild-type TKD are shown. The columns show color-coded Δ%EX of common peptides in the inhibitor complexes at the indicated labeling times. Peptides are assigned peptide ID numbers from the N-terminus (left) to C-terminus (right) on the X-axis. *Supplemental information* lists the amino acid sequence and Δ%EX value of each peptide ID. The position of the secondary structures and conserved HRD and DFG motif regions are shown at the top of the heatmap. The red square box on the heatmaps indicates peptides corresponding to the DFG motif-containing binding pocket. **(C)** The Δ%Ex values of the EGFR TKD with the indicated TKI at 1 min labeling time are color-coded as shown and mapped onto the crystal structure of the EGFR TKD bound to afatinib (4G5J)^47^, dacomitinib (PDB ID:4I23)^57^, and non-covalent osimertinib (See *Results and Discussions* for details). The bound TKI is shown as the stick in each figure. In the crystal structure of the EGFR TKD-dacomitinib complex, the soluble moiety at the R2 position of the inhibitor is not visible; therefore, missing in the structure. All HDX data were obtained from n = 3 separate protein preparations, except for the NC-dacomitinib-bound TKD (n = 2 separate protein preparations), with three independent experiments for each. HDX-MS data statistics are shown in Table S2. The structure maps are generated using PyMOL (http://www.pymol.org).

The overall Δ%Ex patterns of non-covalent (NC) afatinib- or dacomitinib-encounter complex show striking differences despite both inhibitors having the same gefitinib core scaffold (Figures 2B(i-ii), 2C, S11, and S12; Table S2). NC dacomitinib binding stabilizes nearly the entire TKD except for αE: strong engagement with the key functional regions, including the β1×P-loop, αC, the HRD, and DFG motifs are seen (Figures 2B(ii), 2C, S11, and S12). In contrast, the magnitude of the dynamic responses to NC afatinib binding in these regions is much less prominent (Figures 2B(i), 2C, S11, and S12). Most notably, NC afatinib binding does not impact the dynamics of the HRD motif regions and increases αC flexibility in the early labeling time points of 10 sec and 1 min (Figures 2B(i), 2C, S11, and S12). These results demonstrate that similarities in the chemical structure of the inhibitors do not translate into similar protein dynamics.

Another example of inhibitor-dependent dynamics with chemically similar inhibitors is seen with the TKD bound to compound 2 or 3 (Figures S13 and S14; Table S3). Compound 3 has the furan ring of compound 2 at the soluble end replaced with pyrazole, and both compounds were shown to have a superior inhibitory activity to lapatinib by breaking HER2-HER3 heterodimer interaction.^46^ The overall Δ%Ex patterns of the compound 2-or 3-bound TKD show profound differences from that of lapatinib (Figures 1C, 1D, S13, and S14): The overall TKD dynamics with compound 2 is similar to that of erlotinib/gefitinib-bound TKD (Figure 1C-1D) while the compound 3-bound TKD dynamics shows comparable dynamics to a covalent afatinib TKD complex (*See the next result section*).

Where is the possible structural origin of the inhibitor-dependent dynamics? Since the only structural difference between NC afatinib and dacomitinib is chemical substituents at their soluble end (Figure 2A), inhibitor-dependent dynamics must originate from interactions surrounding the soluble end of the inhibitors. An αD region, where the soluble end of these inhibitors interact, is more dynamic in the order of gefitinib (the most flexible) > NC afatinib > NC dacomitinib (the least flexible) at early labeling time points of 10 sec and 1 min (Figure S15A). This finding is consistent with crystal structures bound to these inhibitors^24,47^ where the soluble dimethylamine group of afatinib at the R2 position (Figure 2A) is wedged between D800 of αD and R841 in the binding pocket while the propylmorpholino group of gefitinib points away from these residues and interact with the P-loop (Figure S15B). The dynamics of αD are important not only for propagating inhibitor-dependent dynamics to other regions, as indicated by our HDX-MS but also for facilitating the covalent bond formation with Cys797.^29^ If so, the optimum positioning of the bound inhibitor must involve concerted dynamics between αD and the binding pocket. Because the covalent bond is seen in a crystal structure of the wild-type TKD-afatinib complex solved by inhibitor soaking,^47^ our HDX-MS of the afatinib encounter complex must probe EGFR structures with the bound inhibitor already placed in optimum positions to react. On the other hand, the absence of the covalent bond in a crystal structure of the dacomitinib-TKD solved by soaking suggests the involvement of conformational changes in forming a covalent bond between the inhibitor warhead and Cys797. In other words, a bound dacomitinib must be repositioned to react with Cys797, which requires concerted structural rearrangement between αD and the binding pocket.

If structural plasticity in the encounter complex is the key to forming the covalent bond, a crystal structure of the TKD-inhibitor complex must be solved by co-crystallization. To this end, we solved crystal structures of the wild-type complexed with non-covalent osimertinib by soaking or co-crystallization (Figure S16A; Table S4). A crystal structure comparison between the wild-type TKD bound to NC osimertinib or covalent osimertinib solved by co-crystallization shows significant structural re-arrangements in the N-lobe β-strands and A-loop (Figure S16B). These findings corroborate an observation that a crystal structure of the wild-type TKD with covalent osimertinib has not been solved by crystal soaking,^48^ since crystal packing does not permit the large conformational changes in the N-lobe required for the covalent bond formation. Our HDX-MS result of the osimertinib-encounter complex agrees with these crystal structure comparisons: the entire encounter complex is dynamic except for the β1×P-loop region (Figures 2B(iii), 2C, S11C, and S12). The structural flexibility in the entire TKD should allow the bound osimertinib to explore multiple rotamer conformations for aiding robust covalent bond formation, as suggested by a previous MD simulation study.^48^ Thus, extreme stability of encounter complexes is not a necessary prerequisite to improve the robustness of irreversible inhibitor reactivity.

### A possible stability-specificity trade-off points to a structural origin of TKI side effects

Inhibitor-dependent dynamics in the encounter complexes result in the formation of unique covalent complex structures (Figure 3A, 3B, and S17-S20; Table S5). The αC and binding pocket regions in all covalent complexes are structurally more flexible than those of the corresponding encounter complexes (Figures 3A, 3B, and S18-S20). The β1×P-loop region maintains tight contact with dacomitinib and osimertinib, but not with afatinib (Figures 3A, 3B, and S18-S20). Unlike afatinib and osimertinib complexes, the binding pocket and C-lobe of the dacomitinib complex remain stable (Figures 3A(ii), and S18-S20). Although osimertinib is considered a wild-type sparing inhibitor, thereby lessening side effects compared to the 2^nd^ generation TKIs,^48–50^ it reacts with the wild-type TKD as robustly and efficiently as the 2^nd^ generation TKIs under our reaction condition, as no unreacted wild-type TKD was seen after 5 min incubation with each covalent TKI at pH 7.4, 25°C (Figure S17). Thus, the structural origin of the TKI side effects does not appear to correlate with either the inhibitor reactivity or the dynamics of the covalent complexes, as both afatinib and osimertinb covalent complexes show overall structural similarities except for the β1×P-loop region (Figures 3A and S18-S20). Our HDX-MS results show a better correlation between the severity of side effects and the dynamics of the encounter complexes: the side effects could originate from strong stabilizing effects of the 2^nd^ generation TKI, seen in the dacomitinib encounter complex (Figure 3A(ii), 3B, and S18-S20), by reversibly interacting with off-target proteins as expected from a trade-off between protein-ligand stability and ligand binding specificity.^36^ If so, inhibitors that maximize kinase-inhibitor stability, as sought in conventional inhibitor screenings, may not necessarily produce desirable clinical outcomes with minimum side effects. Our results suggest that a stable interaction between a bound inhibitor and the β1×P-loop region in encounter complexes suffices for the robust formation of the covalent bond, as seen in the non-covalent osimertinib complex (Figures 2B(iii), 2C, S11, and S12). In other words, strong interactions between a bound inhibitor and the binding pocket are not obligatory for targeting kinases with irreversible inhibitors. Our HDX-MS results suggest that the clinical success of osimertinib can be explained by its implicit negative design: The osimertinib encounter complex is structurally dynamic with a disrupted binding pocket (Figures 2B, S11, and S12). Because the ATP binding site is highly conserved among kinases, inhibitors that avoid or disrupt interactions with the binding pocket of encounter complexes may be ideal inhibitors with minimum side effects.

**Figure 3:**
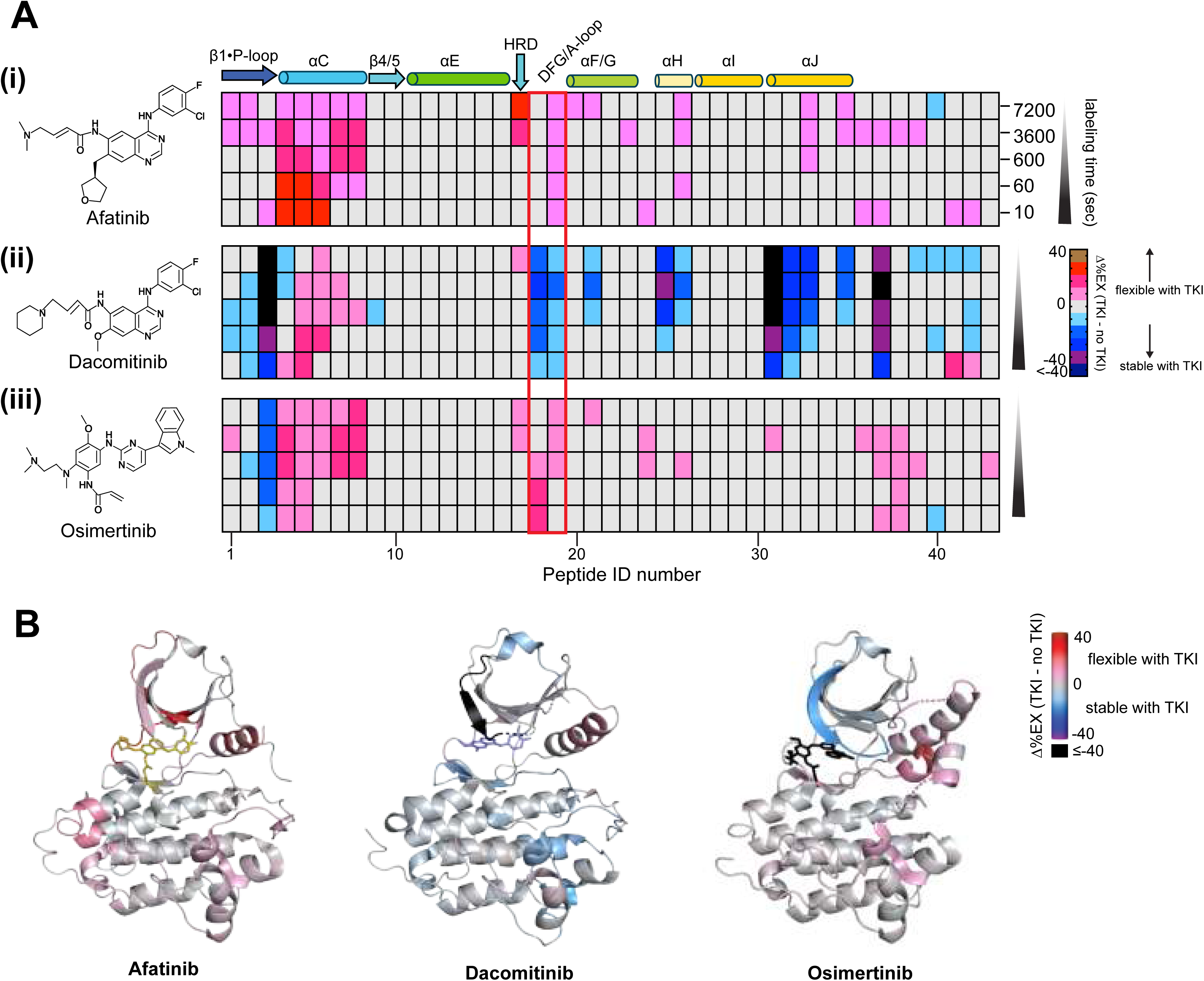
Structural dynamics of the wild-type TKD complexed with the covalent second- and third-generation TKIs probed by HDX-MS. **(A)** The Δ%EX heatmaps of covalent afatinib (i), covalent dacomitinib (ii), or covalent osimertinib (iii)-bound wild-type TKD are shown. The columns show color-coded Δ%EX of common peptides in the inhibitor complexes at the indicated labeling times. Peptides are assigned peptide ID numbers from the N-terminus (left) to C-terminus (right) on the X-axis. *Supplemental information* lists the amino acid sequence and Δ%EX value of each peptide ID. The position of the secondary structures and conserved HRD and DFG motif regions are shown at the top of the heatmap. The red square box on the heatmaps indicates peptides containing the DFG motif and A-loop. **(B)** The Δ%Ex values are mapped onto crystal structures of the wild-type TKD afatinib (PDB ID: 4G5J),^47^ dacomitinib (PDB ID: 4I23),^57^ or osimertinib (PDB ID: 6JXT)^48^ covalent complex at 1 min deuterium labeling time. The bound inhibitor is indicated as a stick in each figure. All HDX data were obtained from n = 3 separate protein preparations, except for the dacomitinib-TKD complex (n = 2 protein preparations), with three independent experiments for each. HDX-MS data statistics are shown in Table S5. The structure maps are generated using PyMOL (http://www.pymol.org).

### Covalent osimertinib causes global stabilization of L858R/T790M

The development of TKIs that specifically target TKI-resistant kinases has been an intense research area.^8,48^ To this end, molecular dynamic simulations have been used to understand how various oncogenic and TKI-resistant mutations influence population distributions of active and inactive kinase structures, thereby affecting TKI binding and selectivity.^27,28^ Typically, in such studies, the “closeness” of simulated results to the canonical active state is evaluated by the degree of αC tilt through the salt bridge distance between K745 in β3 and E762 in αC in addition to A-loop conformations.^27,28^ One such study shows that the most common oncogenic and TKI-resistant EGFR mutations, L858R and L858R/T790M, respectively, increase the population of active EGFR through different mechanisms, the former by destabilizing the inactive conformations and the latter by lowering an energetic barrier to populate conformers with solvent-exposed A-loop.^27,28^ However, T790M is a well-known resistant mutation, even against active-state selective inhibitors, such as gefitinib.^51^ This observation agrees with a view of protein allostery in that no single stable structure consists of a protein ensemble.^43^ If so, an active-state selective inhibitor that targets a protein with certain stable structural features will be rendered ineffective. In other words, structural heterogeneity or flexibility in the T790M ensemble may underlie TKI resistance.

Although HDX-MS is unable to distinguish between the active and inactive conformations based on αC dynamics, unless both conformations are separated by an energetic barrier higher than the thermal energy, both L858R and L858R/T790M mutants in their native states show more flexible αC than that of the wild-type (Figures 4A and 4B(i-ii)). This observation demonstrates that both mutations increase αC’s propensity to deviate from αC dynamics of the wild type, which predominantly populates the inactive state. HDX-MS also shows that these mutations enhance the dynamics of key regions involved in TKI binding, including the hydrophobic pocket, consisting of β4-β5 (Figures 4A and 4B(iii-iv) and the binding pocket containing the conserved HRD and DFG motifs (Figures 4A and 4B(v-vi). The pre-existing flexibility in these regions may prevent an inhibitor from finding its optimum position to prolong its residence time in the binding pocket - a simple explanation for why NSCLC patients carrying these mutations are resistant to reversible first-generation inhibitors despite the administration of active state-selective inhibitors. This observation corroborates the results of kinase activity assays in previous studies where these mutations were shown to cause a decrease in ATP binding affinities.

**Figure 4:**
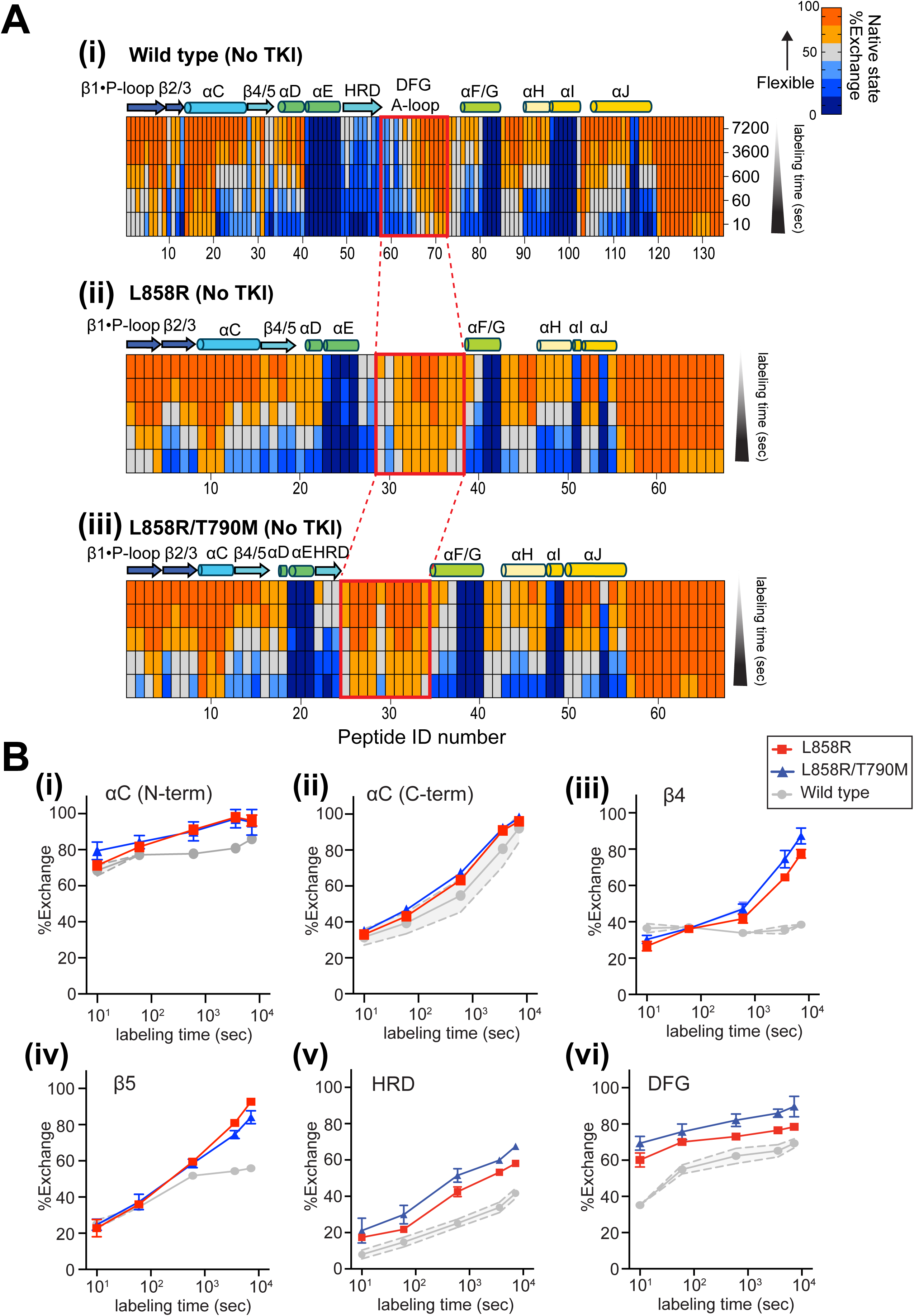
EGFR oncogenic L858R and TKI resistant L858R/T790M mutations increase structural flexibility of the key TKI binding regions. **(A)** The native state percent exchange heatmaps of the wild-type **(i)**, L858R **(ii)**, and L858R/T790M **(iii)**. Each column shows the percent exchange of the same peptides in the wild-type and mutants. Peptides are assigned peptide ID numbers from the N-terminus (left) to C-terminus (right) on the X-axis. *Supplemental information* lists the amino acid sequence and percent exchange value of each peptide ID. The position of the secondary structures and conserved HRD and DFG motif regions are shown at the top of each heatmap. The red square box on the heatmaps indicates peptides corresponding to the DFG motif-containing binding pocket. **(B)** The percent exchange of peptides derived from indicated regions in the wild-type (grey), L858R (red square), and L858R/T790M (blue square) is plotted against the deuterium labeling time (10, 60, 600, 3600, and 7200 sec). The average standard deviation of each peptide in the wild-type TKD is shown as grey error bands. The percent exchange of **(i)** the N-terminal end of αC (residue 752-760), **(ii)** the C-terminal end of αC (residue 767-777), **(iii)** β4 (residue 778-782), **(iv)** β5 (residue 783-788), **(v)** the HRD (residue 827-840), and **(vi)** DFG (residue 845-861) regions. All HDX data were obtained from n = 3 separate protein preparations, with three independent experiments for each. HDX-MS data statistics for L858R and L858R/T790M are presented in Tables S6 and S7.

Osimertinib is the current frontline TKI therapy for NSCLC patients regardless of T790M mutation status.^12–15^ The clinical success of osimertinib is partly explained by its wild-type sparing effect,^52^ though it robustly forms a covalent bond with wild-type (Figure S17). To investigate structural mechanisms of osimertinib clinical success against L858R and L858R/T790M mutants, we probed the dynamics of encounter and covalent complexes of both mutants and compared them to that of the wild type (Figures 5A-5C and S21-S23; Tables S6 and S7). Differences in their dynamics are striking; The encounter and covalent complexes of both wild type and L858R/T790M undergo global dynamic changes, while L858R shows more localized structural changes (Figures 5A-5C, Figures S21-S23). Unlike the wild-type complexes (Figure 5A), destabilized regions in both mutants are localized in αC of L858R/T790M (Figure 5C) and αD of L858R (Figure 5B(ii)). Except for these regions, β-strands in the N-lobe of both mutants show significant structural stabilization (Figures 5B, 5C, Figures S21-S23). Another notable difference between the wild-type and both mutants is found in the DFG-containing binding pocket region, which becomes stable after the covalent bond formation (Figure 5A-5C). The effect of the covalent bond formation is propagated to the C-lobe of the L858R/T790M mutant, resulting in the global stabilization of the entire mutant kinase domain (Figure 5C). Thus, the global stabilization of the L858R/T790M covalent complex, not the encounter complex, may underlie its clinical success.

**Figure 5:**
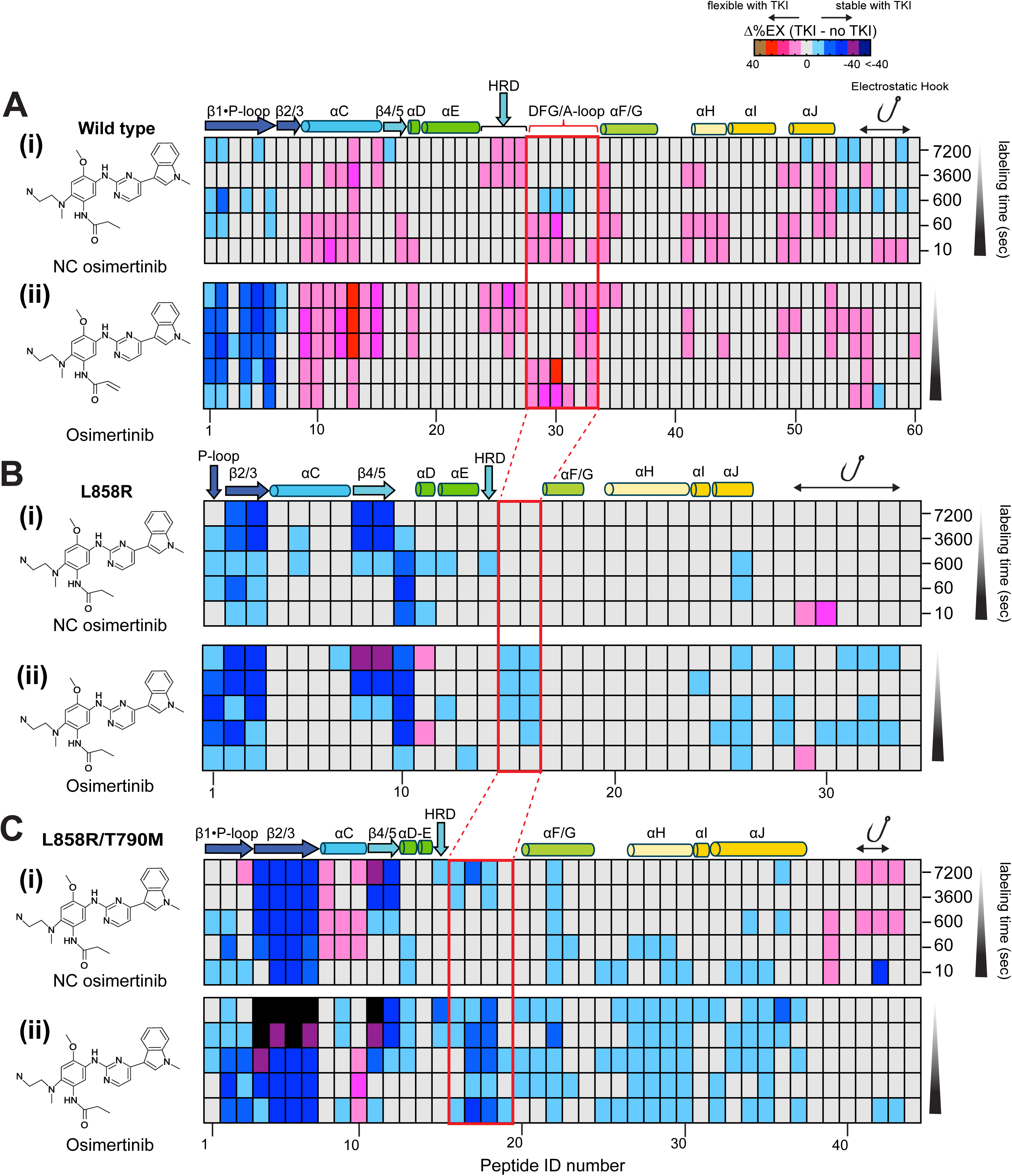
Covalent bond formation with the mutants, but not with the wild-type, stabilizes the TKD. **(A-C)** The Δ%EX heatmaps (TKI - no TKI) of the wild-type **(A)**, L858R **(B)**, and L858R/T790M **(C)** complexed with non-covalent (NC) osimeritnib **(i)** or covalent osimertinib **(ii)**. The columns show color-coded Δ%EX of common peptides in the inhibitor complexes at the indicated labeling times. Peptides are assigned peptide ID numbers from the N-terminus (left) to C-terminus (right) on the X-axis. *Supplemental information* lists the amino acid sequence and Δ%EX value of each peptide ID. The position of the secondary structures and conserved HRD and DFG motif regions are shown at the top of the heatmap. The red square box on the heatmaps indicates peptides in the ATP binding pocket containing the DFG motif and A-loop. All HDX data were obtained from n = 3 separate protein preparations, with three independent experiments for each. HDX-MS data statistics for L858R and L858R/T790M are presented in Tables S6 and S7.

## Conclusions

Our HDX-MS study demonstrates subtle inhibitor-dependent dynamic changes in EGFR TKD-TKI complexes that are seen even with chemically similar inhibitors. Such small structural changes, involving less than 1 Å backbone RMSD, are known to be important for regulating various biological processes, including feedback inhibition, enzyme activation, cooperativity, specificity, and evolutional selection.^53 54,55^ The key driver for these biological processes is protein allostery - proteins’ exquisite ability to sense their subtle structural changes that are propagated to functionally important distant sites. Protein allostery suggests a potential application of negative design to develop drugs that avoid or disrupt interactions with the conserved kinase binding pocket. Such allosteric inhibitors have recently been developed and have shown promising results in a genetically engineered mouse model when another EGFR inhibitor, either cetuximab or osimertinib is co-administered.^20, 56^Allosteric inhibitors, thus, expand the repertoire of combination drug therapy. Studying inhibitor-dependent dynamics in various kinases may further provide new avenues of therapy strategies, such as controlling downstream molecular responses without abolishing kinase activity or restoring wild-type kinase activity levels.

## Method Details

### Cell Culture

#### Insect cells

*Spodoptera frugiperda* Sf9 cells were cultured at 27°C with constant shaking at 110 rpm in serum-free ESF921 Insect Cell Culture Medium (Expression Systems). For the expression of wild type, L858R, or T790M/L858R EGFR TKD, the cells were cultured at 27°C at 90 rpm in the ESF921 medium.

### Protein expression and purification

For the expression of the wild type, L858R, or T790M/L858R TKD, Sf9 cells at 2×10^6^ cells/mL was infected with recombinant baculovirus and cultured for 60-65 h at 27°C. The cells were harvested by centrifugation at 2,200 xg, and the cells were resuspended in cold (4°C) lysis buffer (20 mM Tris/HCl, pH 8.0 containing 500 mM NaCl, 5 mM 2-marcaptoethanol, and 10 %w/v glycerol supplemented with protease inhibitor cocktail (Roche), 50 mL lysis buffer/L cell culture used). Cells were lysed using a microfluidizer (Microfluidics M-110P), and the total cell lysate was centrifuged at 75,000 xg for 1 hour to remove cell debris and insoluble aggregates. The supernatant was filtered through a 0.45 μm filter, and the filtered supernatant was loaded onto a 5 mL HiTrap Chelating Ni^+2^ affinity column (Cytiva) equilibrated in buffer A (20 mM Tris/HCl, pH 7.8 containing 500 mM NaCl). The column was washed using buffer A and 0-20 % step gradient with buffer B (buffer A containing 200 mM imidazole) before an elution step at 100 % buffer B. The pooled fractions were filtered through a 0.22 μm syringe filter, and DTT was added to the final concentration of 10 mM. The sample was loaded onto HiLoad 16/60 Superdex 200 column (Cytiva) equilibrated in 20 mM HEPES, 500 mM NaCl at pH 7.4). The purity and molecular weight of wild type, L858R, or T790M/L858R EGFR TKD were assessed and verified by SDS-PAGE and mass spectrometry. The purified TKD in size exclusion fractions was used for HDX-MS without concentration.

### Hydrogen-Deuterium Exchange and Mass Spectrometry (HDX-MS)

Deuterium-labeled samples were prepared immediately after protein purification by LEAP HDX-2 automation system (Trajan) or manual labeling. All L858R and L858R/T790M mutant data were collected using manually labeled samples. The automation system prepared deuterium-labeled wild type TKD by pipetting 4 μL of 0.3 mg/mL protein (7.8 μM) at 4°C and mixing 76 μL of the deuterium buffer (20 mM HEPES, 100 mM NaCl, pD 7.4) at 25°C. The deuterium labeling was quenched at different time points by the addition of cold (4°C) quench buffer (80 μL of 200 mM glycine, 0.1 M TCEP, pH 2.4). The labeled protein sample (100 μL) was immediately injected onto a pepsin column (Waters) maintained at 2°C in the HDX manager (Waters) and digested for 3 min while flowing water (Honeywell Burdick and Jackson) with 0.1 % (v/v) formic acid (solvent A) at 100 μL/min. The peptic peptides were trapped in BEH C18 column (2.1 x 5 mm, 1.7 μm, Waters) equilibrated in 5 % solvent B (acetonitrile (Honeywell Burdick and Jackson) with 0.1 % (v/v) formic acid) and separated using Acquity UPLC BEH C18 column (1.0 × 100 mm, 1.7 μm, 130 Å, Waters) at 2°C. The peptic peptides were eluted using a linear gradient from 5 to 40 % solvent B over 7 min at 40 μL/min and analyzed using a Synapt G2-Si mass spectrometer (Waters) by acquiring MS^e^ data. The data were collected in the sensitivity TOF mode at 0.4 s scan time with low and high collision energy of 5 to 10 V and 15 to 40 V, respectively, with the scan range between 300 and 1500 m/z. Other instrument parameters are the followings: 3.0 kV capillary voltage, 20 V cone voltage, 80 V source offset, 100°C source temperature, and 150°C desolvation temperature, 600 L/h desolvation gas flow, and 6 bar nebulizer gas. To prepare deuterium-labeled wild type TKD with various inhibitors, the TKD (0.3 mg/mL – 0.6 mg/mL, 7.8 μM – 15.6 μM) was pre-incubated with erlotinib (15.6 to 20 μM), gefitinib (20 μM), lapatinib (20 μM), non-covalent afatinib or dacomitinib (20 μM), or non-covalent osimertinib (30 μM) at 25°C for 30 min prior to the deuterium labeling. To prepare a covalent TKD complex, the TKD at 0.6 mg/mL (15.6 μM) was incubated with 20 μM of covalent afatinib, dacomitinib, or osimertinib at 25°C for 5 min to 1 hour, and the covalent complex formation was verified by intact mass analysis using mass spectrometry (*See Figure S17*) prior to the deuterium labeling experiments. To prepare manually labeled HDX-MS samples, 5 μL L858R or T790M/L858R at 0.3-0.6 mg/mL (7.8 μM – 15.6 μM) was mixed with 95 μL of deuterium buffer (20 mM HEPES, 100 mM NaCl, pD 7.4) at 25°C. The labeling was quenched at different time points (10, 60, 600, 3600, or 7200 sec) by the addition of 100 μL cold quench buffer (20 mM glycine, 0.1 M TCEP, pH 2.4), and the labeled samples were immediately flash-frozen in liquid N_2_. The frozen samples were stored at -80°C until mass spectrometry analysis. To prepare no deuterium or full-deuterated deuterium labeled protein samples, 20 mM HEPES, 100 mM NaCl at pH 7.4 or 20 mM HEPES, 100 mM NaCl, 8 M urea-d4 (Cambridge Isotope Labs) at pD 7.4, respectively, was used instead of using the standard deuterium labeling buffer (20 mM HEPES, 100 mM NaCl, pD 7.4).

### HDX-MS data analysis

Peptic peptides were sequenced using the ProteinLynx Global Server 3.03 (PLGS, Waters), and the deuterium uptake of each peptide was assessed using DynamX 3.0 (Waters) with the intensity cut-off at 5,000, minimum product per amino acid to 0.2-0.3, and maximum MH^+^ error at 10 ppm. The deuterium uptake of all peptides reported in this study is the average uptake of three biological samples (two for the wild type TKD with non-covalent dacomitinib) performed in technical triplicates. The percent exchange of each peptide (%Ex) was calculated by the following equation:

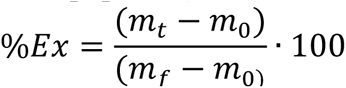

where m_t_ = the centroid mass of a peptic peptide at time, t, m_0_ = the centroid mass of a peptic peptide without deuterium labeling, and m_f_ = the centroid mass of a peptic peptide for the fully-deuterated standard protein. The Pymol script files to prepare the Δ%Ex structure maps in Figure 2,4-6, Figure S2, S4, S7, S11, S14, and S15 were made using DynamX 3.0 (Waters). Briefly, to assign %Ex of single residues, %Ex of the shortest peptide in a given region was used over longer peptides containing the same region. If multiple overlapping peptides with the same length were present for a given region, %Ex of a peptide covering more C-terminal end of a region was used. All data were collected and analyzed according to consensus HDX-MS guidelines.^58^ Statistical analysis of all HDX-MS data presented in Tables S1-S3 and S5-S7 was performed as described previously.^59^

### Intact mass analysis

The purified wild type and mutant TKD (0.3-0.6 mg/mL) was incubated with a covalent inhibitor at 1:2 (protein:TKI) molar ratio at 25°C in 20 mM HEPES, 100 mM NaCl at pH 7.4. The formation of covalent TKI-TKD complex was monitored by determining the mass of the covalent complex at different time points (Figure S9). An incubated sample was injected onto a mass prep column (Waters) equilibrated in 50:50 H_2_O:acetonitrile containing 0.1 % formic acid followed by the protein elution using a step gradient from 5 % to 98 % acetonitrile over 5 min. MS spectra of the protein sample was acquired on Waters Xevo-QTOF or Synapt G2-Si by scanning 300-2000 m/z at 0.4 /sec scan rate. The MS spectra was deconvoluted using MassLynx MaxEnt1 (Waters) to determine the mass of the TKD-covalent TKI complex.

### Crystal structure determination of wild-type EGFR TKD complexed with non-covalent osimertinib by soaking or co-crystallization

Purified EGFR TKD (residue 696-1022) was subjected to an additional anion exchange chromatography step, concentrated to ∼5 mg/ml in 20 mM Tris pH 7.5-8.0, 250 mM NaCl and 2.5% 1,3-propanediol. For co-crystallization, a 1:1.5 molar ratio of protein and inhibitor was prepared using a 10 mM stock solution of non-covalent osimertinib in DMSO. The corresponding solution was incubated at 4 °C (using ice bath) for 15 min and then centrifuged at 10,000 *g* for 5 min prior to the crystallization. Crystals were successfully obtained at 16 °C using the hanging-drop vapor diffusion method with a 1:1 volume ratio of reservoir: protein solution; the reservoir solution constituted of 100 mM MES, pH 6.0, 0.8 M sodium citrate. Crystals appeared after 3-4 weeks and were cryo-protected in the reservoir solution supplemented with 20% (w/v) glycerol and 5% (w/v) ethylene glycol prior to flash freezing in liquid nitrogen. For a soaking experiment, the wild-type protein at ∼ 5.5 mg/ml was employed; crystals appeared after 2-3 weeks in a reservoir solution of 100 mM HEPES, pH 7.0, 0.6-0.9 Na-K tartrate. Crystals of the apo-kinase domain appeared after 2-3 weeks. A stock solution of ∼1 mM non-covalent osimertinib was prepared using 200 mM HEPES, pH 7.0, 1.5 M Na-K tartrate. This solution was filtered through a 0.2 μm mesh filter to remove insoluble particulates prior to use. Using this stock solution, approximately 2 molar excess of the inhibitor was introduced to the corresponding hanging drops containing the crystals of EGFR TKD, and the lid was closed for incubation over 1 h at 16 °C. The crystals were then harvested and cryo-protected in the reservoir solution supplemented with 20% (w/v) glycerol and 5% (w/v) ethylene glycol prior to flash freezing in liquid nitrogen.

X-ray diffraction data were collected on the synchrotron beamline 24-ID-E of NE-CAT at the Advanced Photon Source. Data were processed using XDS Version 20200417 and scaled using SCALA (Version 3.3.22) in the CCP4 program suite (Version 7.1).^60^ The structures were solved by molecular replacement using Phaser.^61^ The active state EGFR TKD structure (PDB code 1M17)^23^ was used as the search model. Repeated cycles of manual building/rebuilding were performed using Coot^62^ and were alternated with refinements using Phenix (Version 1.14_3260)^63^ and the PDB-REDO web server (https://pdb-redo.eu).^64^ The final structure was validated with the MolProbity (version 4.02b-467)^65^ and wwPDB (version 2.26) servers. Structural figures were generated using PyMOL (http://www.pymol.org).

### Synthesis of non-covalent afatinib, dacomitinib, and osimertinib

All reactions were carried out in flame-dried glassware under a vacuum and then purged and cooled under an atmosphere of dry nitrogen or argon. Molecular sieves (4Å) were dried at 160 °C under 0.25 mtorr vacuum prior to use. Unless otherwise mentioned, solvents were purified as follows. Methanol and 1,2-dimethoxyethane were purchased from Sigma Aldrich and incubated over 4Å MS for 48 h prior to use. CH_2_Cl_2_ was dried over CaH_2_ and freshly distilled prior to use. NMR spectra were obtained using a 400 MHz Bruker NMR spectrometer and referenced using the residual ^1^H peak from the deuterated solvent. Agilent 6550A (QTOF) instrument was used for HRMS (ESI) analysis with polyethylene glycol (PEG-400-600) as a reference. Column chromatography was performed using Silicycle 60Å, 35-75 µm silica gel. Pre-coated 0.25 mm thick silica gel 60 F254 plates were used for analytical thin layer chromatography (TLC) and visualized using UV light, iodine, potassium permanganate stain, *p*-anisaldehyde stain or phosphomolybdic acid in EtOH stain. All other commercially available reagents and solvents were used as received unless otherwise mentioned.

### Synthesis of non-covalent osimertinib

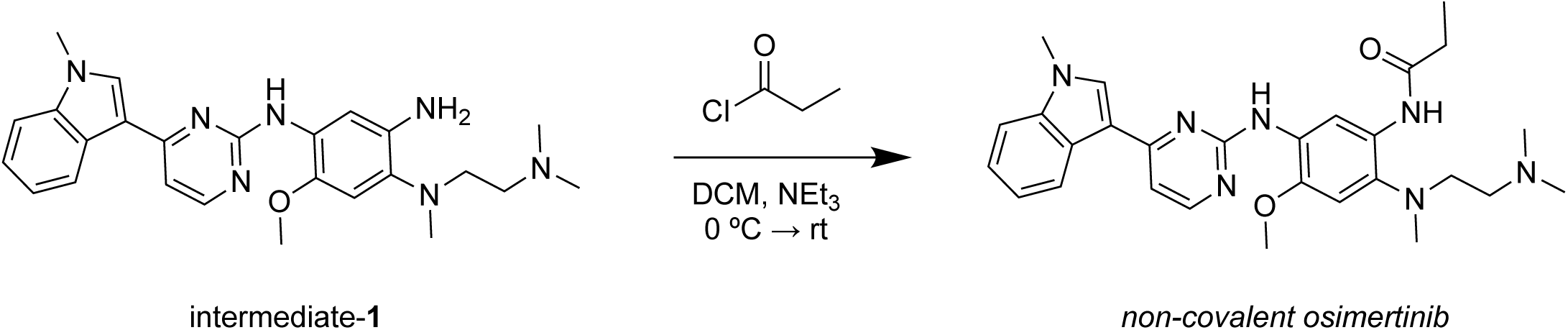

#### Synthesis of *N*-(2-((2-(dimethylamino)ethyl)(methyl)amino)-4-methoxy-5-((4-(1-methyl-1*H*-indol-3-yl)pyrimidin-2-yl)amino)phenyl)propionamide (non-covalent osimertinib)

Intermediate-1 (MedChemExpress; HY-78869) obtained commercially, was repurified via basic silica gel chromatography using a mixture of MeOH/DCM supplemented with 1% NH_4_OH. 100 mg of purified intermediate-1 (0.224 mmol, 1.0 equiv) was dissolved in DCM to a final concentration of 0.1 M in a round bottom flask equipped with a magnetic stirrer. To this solution, 94 μL of triethylamine (0.673 mmol, 3.0 equiv) was added, and the solution was cooled to about 0 °C using an ice-water bath. To this ice-cold solution, a 0.1 M solution of propionyl chloride (0.269 mmol, 1.2 equiv) in DCM was added dropwise over a period of 1 h. The reaction mixture was then allowed to reach room temperature and stirred further for another hour. The progress of this reaction was monitored by TLC. The mixture was then poured over ice-cold solution of 10% NaHCO_3_ in a separatory funnel and the DCM layer was extracted, washed twice with 10% NaHCO_3_, brine and dried over anhydrous Na_2_SO_4_. DCM was then removed by rotaryvapor and the resulting crude product was purified via basic silica gel chromatography.

#### Analytical data for *N*-(2-((2-(dimethylamino)ethyl)(methyl)amino)-4-methoxy-5-((4-(1-methyl-1*H*-indol-3-yl)pyrimidin-2-yl)amino)phenyl)propionamide

The purified product was obtained as an off-white amorphous powder in about 42% overall yield. ^1^H NMR (400 MHz, CDCl_3_) δ 1.30 (t, 3H, *J* = 7.6 Hz), 2.27-2.31 (m, 8H), 2.46 (dd, 2H, *J* = 7.6 Hz), 2.68 (s, 3H), 2.92 (t, 2H, *J* = 6.0 Hz), 3.78 (s, 3H), 4.00 (s, 3H), 6.78 (s, 1H), 7.20 (d, 1H, *J* = 6.0 Hz), 7.25-7.28 (m, 2H), 7.4 (dt, 1H, *J* = 6.5, 3.0 and 1.0 Hz), 7.72 (s, 1H,), 8.07 (dd, 1H, *J* = 6.5 and 3.0 Hz), 8.38 (d, 1H, *J* = 5.2 Hz), 9.09 (s, broad, 1H), 9.69 (s, broad, 1H), 9.74 (s, 1H); ^13^C NMR (CDCl_3_, 100 MHz) δ 10.3, 33.2, 44.2, 45.6, 56.4, 57.6, 104.8, 108.1, 109.9, 110.2, 113.9, 120.5, 121.1, 121.9, 126.2, 127.9, 130.0, 134.2, 135.3, 138.5, 144.1, 158.1, 159.8, 162.3, 171.5. HRMS: ES^+^ (C_28_H_35_N_7_O_2_): Calc. [M + H]^+^: 502.2930, Found [M + H]^+^: 502.2926 (Figures S8-S9).

### Synthesis of non-covalent afatinib

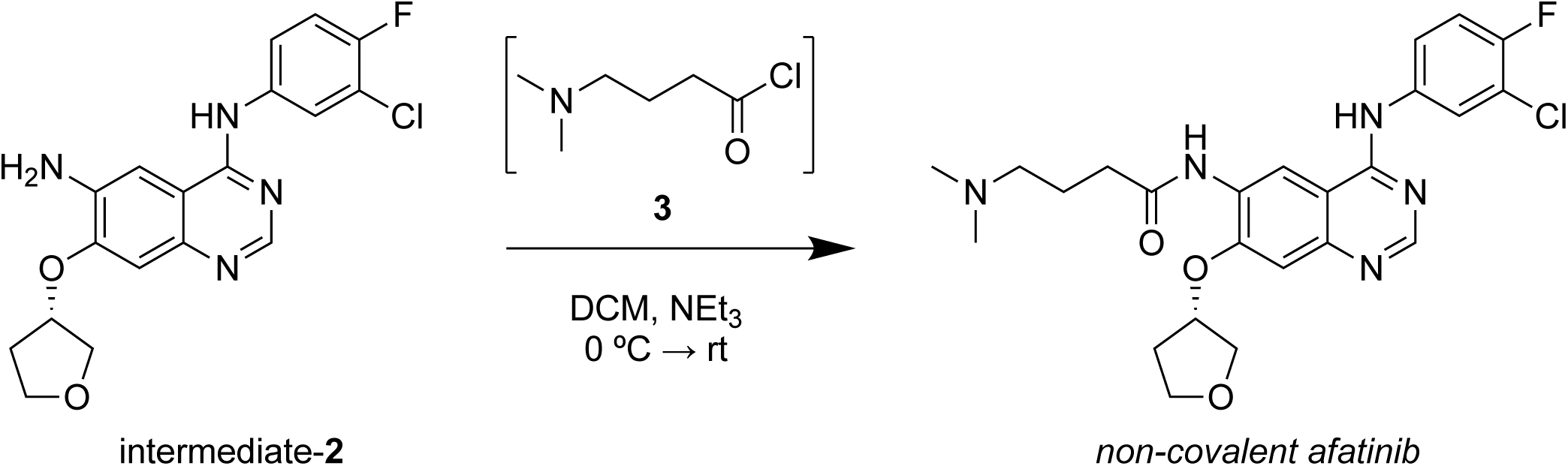

#### Synthesis of *(S)*-N-(4-((3-chloro-4-fluorophenyl)amino)-7-((tetrahydrofuran-3-yl)oxy) quinazolin-6-yl)-4-(dimethylamino)butanamide (non-covalent afatinib)

4-dimethylaminobutyric acid hydrochloride (54 mg, 0.32 mmol, 1.2 equiv, Combi-Blocks, QC-1214) was transferred to a pear-shaped flask containing 3 mL of DCM. The slurry was stirred and cooled to 0 °C using an ice-water bath. 47 μL of thionyl chloride (2.4 equiv) was added dropwise and the mixture was warmed to room temperature and stirred overnight. The volatile contents were removed under a stream of dry nitrogen and later under vacuum. The resulting crude residue (**3**) was suspended in 2 mL of DCM and used directly for the next step without further purification. Commercially procured (Ambeed; A218083) intermediate-2 (100 mg, 0.267 mmol, 1.0 equiv) was dissolved in DCM to a final concentration of 0.1 M in a round bottom flask equipped with a magnetic stirrer. To this solution 110 μL of triethylamine (0.8 mmol, 3.0 equiv) were added and the solution was cooled to about 0 °C using an ice-water bath. To this ice-cold solution, the above solution of crude 4-dimethylaminobutanoyl chloride (**3**) was added dropwise over a period of 1 h. The reaction mixture was then allowed to reach room temperature and stirred further for another hour. The progress of this reaction was monitored by TLC. The mixture was then poured over ice-cold solution of 10% NaHCO_3_ in a separatory funnel and the DCM layer was extracted, washed twice with 10% NaHCO_3_, brine and dried over anhydrous Na_2_SO_4_. DCM was then removed by rotaryvapor and the resulting crude product was purified via basic silica gel chromatography.

#### Analytical data for *(S)*-N-(4-((3-chloro-4-fluorophenyl)amino)-7-((tetrahydrofuran-3-yl)oxy)quinazolin-6-yl)-4-(dimethylamino)butanamide

The purified product was obtained as a white amorphous powder in about 78% overall yield. ^1^H NMR (400 MHz, CDCl_3_) δ 1.93-2.0 (m, 3H), 2.17-2.23 (m, 1H), 2.31-2.38 (m, 7H), 2.55-2.59 (m, 4H), 2.99 (dd, 1H, *J* = 14.4 and 7.2 Hz), 3.88 (dt, 1H, *J* = 8.5 and 5.2 Hz), 3.97-4.10 (m, 3H), 5.04 (dd, 1H, *J* = 4.5 and 5.6 Hz), 7.02-7.07 (m, 2H,), 7.45 (m, 1H), 7.77 (dd, 1H, *J* = 6.5 and 2 Hz), 8.05 (s, broad, 1H), 8.44 (s, broad, 1H), 8.56 (s, 1H), 8.97 (s, broad, 1H); ^13^C NMR (CDCl_3_, 100 MHz) δ 9.0, 22.3, 22.7, 33.0, 35.0, 44.6, 45.9, 57.9, 67.5, 73.1, 79.5, 108.5, 109.5, 111.6, 116.5 (d, *J* = 22), 120.8 (d, *J* = 18),122 (d, *J* = 6.5), 124.3, 128.0, 135.5 (d, *J* = 3.5), 149.9 (d, *J* = 305), 154.8 (d, *J* = 244.4), 155.8 (d, *J* = 234.3), 171.8, 176.3.

HRMS: ES^+^ (C_24_H_27_ClFN_5_O_3_): Calc. [M + H]^+^: 488.1865, Found [M + H]^+^: 488.1870 (Figure S4-S5).

### Synthesis of non-covalent dacomitinib

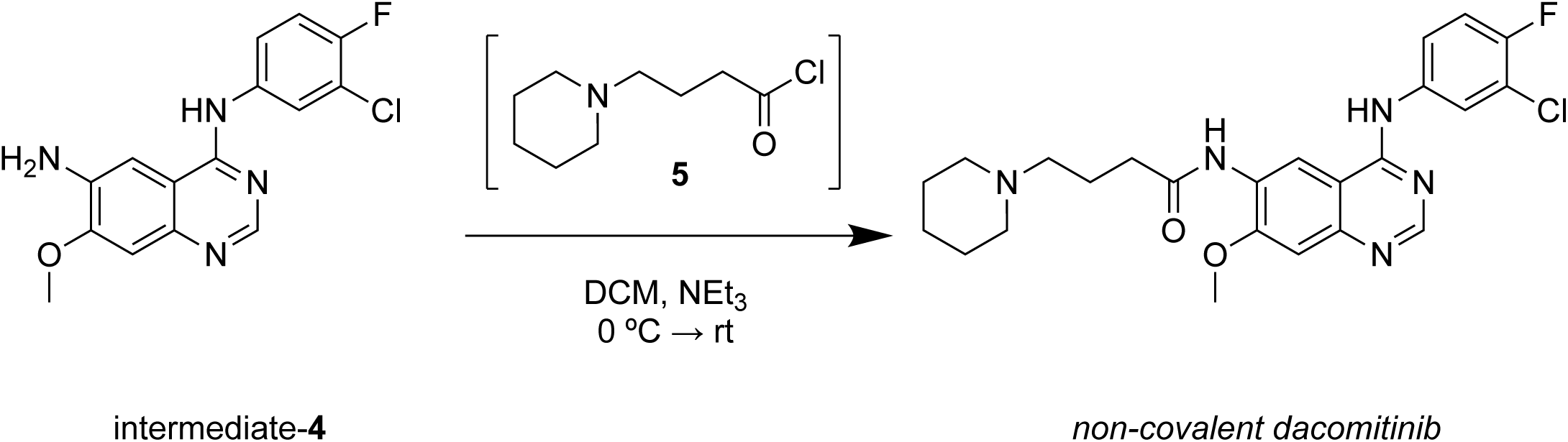

#### Synthesis of N-(4-((3-chloro-4-fluorophenyl)amino)-7-methoxyquinazolin-6-yl)-4-(piperidin-1-yl)butanamide (non-covalent dacomitinib)

4-(piperidin-1-yl)butanoic acid hydrochloride (78 mg, 0.38 mmol, 1.2 equiv, AstaTech, G29935) was transferred to a pear-shaped flask containing 3 mL of DCM. The slurry was stirred and cooled to 0 °C using an ice-water bath. Thionyl chloride (55 μL, 2.4 equiv) was added dropwise, and the mixture was warmed to room temperature and stirred overnight. The volatile contents were removed under a stream of dry nitrogen and later under vacuum. The resulting crude slurry of **5** was suspended in 2 mL of DCM and used directly for the next step without further purification. Commercially procured (Ambeed; A200880) intermediate-4 (100 mg, 0.314 mmol, 1.0 equiv) was dissolved in DCM to a final concentration of 0.1 M in a round bottom flask equipped with a magnetic stirrer. To this solution, 131 μL of triethylamine (0.94 mmol, 3.0 equiv) was added, and the solution was cooled to about 0 °C using an ice-water bath. To this ice-cold solution, the above solution of 4-(piperidin-1-yl)butanoyl chloride was added dropwise over a period of 1 h. The reaction mixture was then allowed to reach room temperature and stirred further for another hour. The progress of this reaction was monitored by TLC. The mixture was then poured over ice-cold solution of 10% NaHCO_3_ in a separatory funnel and the DCM layer was extracted, washed twice with 10% NaHCO_3_, brine and dried over anhydrous Na_2_SO_4_. DCM was then removed by rotaryvapor and the resulting crude product was purified via basic silica gel chromatography.

### Analytical data for N-(4-((3-chloro-4-fluorophenyl)amino)-7-methoxyquinazolin-6-yl)-4-(piperidin-1-yl)butanamide

The purified product was obtained as a white amorphous powder in about 61% overall yield. As seen from the NMR spectrum, traces of methanol (H-bonded to amide N-H) were observed. ^1^H NMR (400 MHz, CDCl_3_) δ 1.44 (m, 2H), 1.56 (m, 4H), 1.93 (dd, 2H, *J* = 13.6 and 6.8 Hz), 2.39 (m, 7H), 2.52 (t, 2H, *J* = 7.2 Hz), 4.02 (s, 3H), 7.10 (t, 1H, *J* = 8.4 Hz), 7.21 (s, 1H), 7.49 (m, 1H), 7.80 (d, 1H, *J* = 0.5 Hz), 7.86 (dd, 1H, *J* = 2.4 and 1.2 Hz), 8.41 (s, broad, 1H), 8.62 (s, 1H), 9.00 (s, broad, 1H); ^13^C NMR (CDCl_3_, 100 MHz) δ 22.8, 24.6, 26.1, 36.0, 39.2, 54.6, 56.4, 57.8, 106.7, 106.9, 109.6 (d, *J* = 42.8), 116.5 (d, *J* = 22), 121.0 (d, *J* = 18.4), 121.7 (d, *J* = 6.7), 124.2, 128.1, 135.4 (d, *J* = 3.1), 149.1 (d, *J* = 141), 153.0, 154.8 (d, *J* = 244.3), 155.7 (d, *J* = 238.4), 172.4. HRMS: ES^+^ (C_24_H_27_ClFN_5_O_3_): Calc. [M + H]^+^: 472.1916, Found [M + H]^+^: 472.1922 (Figures S6-S7).

## References

1. Freudlsperger, C., Burnett, J.R., Friedman, J.A., Kannabiran, V.R., Chen, Z., and Van Waes, C. (2011). EGFR-PI3K-AKT-mTOR signaling in head and neck squamous cell carcinomas: attractive targets for molecular-oriented therapy. Expert Opin Ther Targets 15, 63–74. 10.1517/14728222.2011.541440.

2. Lurje, G., and Lenz, H.J. (2009). EGFR signaling and drug discovery. Oncology 77, 400–410. 10.1159/000279388.

3. Molina, J.R., and Adjei, A.A. (2006). The Ras/Raf/MAPK pathway. J Thorac Oncol 1, 7–9.

4. Melosky, B., Kambartel, K., Hantschel, M., Bennetts, M., Nickens, D.J., Brinkmann, J., Kayser, A., Moran, M., and Cappuzzo, F. (2022). Worldwide Prevalence of Epidermal Growth Factor Receptor Mutations in Non-Small Cell Lung Cancer: A Meta-Analysis. Mol Diagn Ther 26, 7–18. 10.1007/s40291-021-00563-1.

5. Stewart, E.L., Tan, S.Z., Liu, G., and Tsao, M.S. (2015). Known and putative mechanisms of resistance to EGFR targeted therapies in NSCLC patients with EGFR mutations-a review. Transl Lung Cancer Res 4, 67–81. 10.3978/j.issn.2218-6751.2014.11.06.

6. Wood, E.R., Truesdale, A.T., McDonald, O.B., Yuan, D., Hassell, A., Dickerson, S.H., Ellis, B., Pennisi, C., Horne, E., Lackey, K., et al. (2004). A unique structure for epidermal growth factor receptor bound to GW572016 (Lapatinib): relationships among protein conformation, inhibitor off-rate, and receptor activity in tumor cells. Cancer Res 64, 6652–6659. 10.1158/0008-5472.CAN-04-1168.

7. Opdam, F.L., Guchelaar, H.J., Beijnen, J.H., and Schellens, J.H. (2012). Lapatinib for advanced or metastatic breast cancer. Oncologist 17, 536–542. 10.1634/theoncologist.2011-0461.

8. Du, X., Yang, B., An, Q., Assaraf, Y.G., Cao, X., and Xia, J. (2021). Acquired resistance to third-generation EGFR-TKIs and emerging next-generation EGFR inhibitors. Innovation (Camb) 2, 100103. 10.1016/j.xinn.2021.100103.

9. Sullivan, I., and Planchard, D. (2016). Next-Generation EGFR Tyrosine Kinase Inhibitors for Treating EGFR-Mutant Lung Cancer beyond First Line. Front Med (Lausanne) 3, 76. 10.3389/fmed.2016.00076.

10. Dacomitinib Approved, but Might Not Be Used. (2018). Cancer Discov 8, 1500. 10.1158/2159-8290.CD-NB2018-138.

11. Shyam Sunder, S., Sharma, U.C., and Pokharel, S. (2023). Adverse effects of tyrosine kinase inhibitors in cancer therapy: pathophysiology, mechanisms and clinical management. Signal Transduct Target Ther 8, 262. 10.1038/s41392-023-01469-6.

12. Finlay, M.R., Anderton, M., Ashton, S., Ballard, P., Bethel, P.A., Box, M.R., Bradbury, R.H., Brown, S.J., Butterworth, S., Campbell, A., et al. (2014). Discovery of a potent and selective EGFR inhibitor (AZD9291) of both sensitizing and T790M resistance mutations that spares the wild type form of the receptor. J Med Chem 57, 8249–8267. 10.1021/jm500973a.

13. Reck, M., and Rabe, K.F. (2017). Precision Diagnosis and Treatment for Advanced Non-Small-Cell Lung Cancer. N Engl J Med 377, 849–861. 10.1056/NEJMra1703413.

14. Ramalingam, S.S., Vansteenkiste, J., Planchard, D., Cho, B.C., Gray, J.E., Ohe, Y., Zhou, C., Reungwetwattana, T., Cheng, Y., Chewaskulyong, B., et al. (2020). Overall Survival with Osimertinib in Untreated, EGFR-Mutated Advanced NSCLC. N Engl J Med 382, 41–50. 10.1056/NEJMoa1913662.

15. Passaro, A., Mok, T.S.K., Attili, I., Wu, Y.L., Tsuboi, M., de Marinis, F., and Peters, S. (2023). Adjuvant Treatments for Surgically Resected Non-Small Cell Lung Cancer Harboring EGFR Mutations: A Review. JAMA Oncol 9, 1124–1131. 10.1001/jamaoncol.2023.0459.

16. Thress, K.S., Paweletz, C.P., Felip, E., Cho, B.C., Stetson, D., Dougherty, B., Lai, Z., Markovets, A., Vivancos, A., Kuang, Y., et al. (2015). Acquired EGFR C797S mutation mediates resistance to AZD9291 in non-small cell lung cancer harboring EGFR T790M. Nat Med 21, 560–562. 10.1038/nm.3854.

17. Tumbrink, H.L., Heimsoeth, A., and Sos, M.L. (2021). The next tier of EGFR resistance mutations in lung cancer. Oncogene 40, 1–11. 10.1038/s41388-020-01510-w.

18. Mansour, M.A., AboulMagd, A.M., Abbas, S.H., Abdel-Rahman, H.M., and Abdel-Aziz, M. (2023). Insights into fourth generation selective inhibitors of (C797S) EGFR mutation combating non-small cell lung cancer resistance: a critical review. RSC Adv 13, 18825–18853. 10.1039/d3ra02347h.

19. Shaikh, M., Shinde, Y., Pawara, R., Noolvi, M., Surana, S., Ahmad, I., and Patel, H. (2022). Emerging Approaches to Overcome Acquired Drug Resistance Obstacles to Osimertinib in Non-Small-Cell Lung Cancer. J Med Chem 65, 1008–1046. 10.1021/acs.jmedchem.1c00876.

20. Marasco, M., and Misale, S. (2022). Resistance is futile with fourth-generation EGFR inhibitors. Nat Cancer 3, 381–383. 10.1038/s43018-022-00365-2.

21. Arter, C., Trask, L., Ward, S., Yeoh, S., and Bayliss, R. (2022). Structural features of the protein kinase domain and targeted binding by small-molecule inhibitors. J Biol Chem 298, 102247. 10.1016/j.jbc.2022.102247.

22. Haddad, Y., Remes, M., Adam, V., and Heger, Z. (2021). Toward structure-based drug design against the epidermal growth factor receptor (EGFR). Drug Discov Today 26, 289–295. 10.1016/j.drudis.2020.10.007.

23. Stamos, J., Sliwkowski, M.X., and Eigenbrot, C. (2002). Structure of the epidermal growth factor receptor kinase domain alone and in complex with a 4-anilinoquinazoline inhibitor. J Biol Chem 277, 46265–46272. 10.1074/jbc.M207135200.

24. Yun, C.H., Boggon, T.J., Li, Y., Woo, M.S., Greulich, H., Meyerson, M., and Eck, M.J. (2007). Structures of lung cancer-derived EGFR mutants and inhibitor complexes: mechanism of activation and insights into differential inhibitor sensitivity. Cancer Cell 11, 217–227. 10.1016/j.ccr.2006.12.017.

25. Park, J.H., Liu, Y., Lemmon, M.A., and Radhakrishnan, R. (2012). Erlotinib binds both inactive and active conformations of the EGFR tyrosine kinase domain. Biochem J 448, 417–423. 10.1042/BJ20121513.

26. Roskoski, R., Jr. (2016). Classification of small molecule protein kinase inhibitors based upon the structures of their drug-enzyme complexes. Pharmacol Res 103, 26–48. 10.1016/j.phrs.2015.10.021.

27. Sutto, L., and Gervasio, F.L. (2013). Effects of oncogenic mutations on the conformational free-energy landscape of EGFR kinase. Proc Natl Acad Sci U S A 110, 10616–10621. 10.1073/pnas.1221953110.

28. Galdadas, I., Carlino, L., Ward, R.A., Hughes, S.J., Haider, S., and Gervasio, F.L. (2021). Structural basis of the effect of activating mutations on the EGF receptor. Elife 10. 10.7554/eLife.65824.

29. Capoferri, L., Lodola, A., Rivara, S., and Mor, M. (2015). Quantum mechanics/molecular mechanics modeling of covalent addition between EGFR-cysteine 797 and N-(4-anilinoquinazolin-6-yl) acrylamide. J Chem Inf Model 55, 589–599. 10.1021/ci500720e.

30. Gao, J., Jian, J., Jiang, Z., and Van Schepdael, A. (2023). Screening assays for tyrosine kinase inhibitors: A review. J Pharm Biomed Anal 223, 115166. 10.1016/j.jpba.2022.115166.

31. Gao, K., Oerlemans, R., and Groves, M.R. (2020). Theory and applications of differential scanning fluorimetry in early-stage drug discovery. Biophys Rev 12, 85–104. 10.1007/s12551-020-00619-2.

32. Kitchen, D.B., Decornez, H., Furr, J.R., and Bajorath, J. (2004). Docking and scoring in virtual screening for drug discovery: methods and applications. Nat Rev Drug Discov 3, 935–949. 10.1038/nrd1549.

33. Batool, M., Ahmad, B., and Choi, S. (2019). A Structure-Based Drug Discovery Paradigm. Int J Mol Sci 20. 10.3390/ijms20112783.

34. Gagic, Z., Ruzic, D., Djokovic, N., Djikic, T., and Nikolic, K. (2019). In silico Methods for Design of Kinase Inhibitors as Anticancer Drugs. Front Chem 7, 873. 10.3389/fchem.2019.00873.

35. Havranek, J.J., and Harbury, P.B. (2003). Automated design of specificity in molecular recognition. Nat Struct Biol 10, 45–52. 10.1038/nsb877.

36. Bolon, D.N., Grant, R.A., Baker, T.A., and Sauer, R.T. (2005). Specificity versus stability in computational protein design. Proc Natl Acad Sci U S A 102, 12724–12729. 10.1073/pnas.0506124102.

37. Karachaliou, N., Fernandez-Bruno, M., Bracht, J.W.P., and Rosell, R. (2019). EGFR first- and second-generation TKIs-there is still place for them in EGFR-mutant NSCLC patients. Transl Cancer Res 8, S23–S47. 10.21037/tcr.2018.10.06.

38. van Alderwerelt van Rosenburgh, I.K., Lu, D.M., Grant, M.J., Stayrook, S.E., Phadke, M., Walther, Z., Goldberg, S.B., Politi, K., Lemmon, M.A., Ashtekar, K.D., and Tsutsui, Y. (2022). Biochemical and structural basis for differential inhibitor sensitivity of EGFR with distinct exon 19 mutations. Nat Commun 13, 6791. 10.1038/s41467-022-34398-z.

39. James, E.I., Murphree, T.A., Vorauer, C., Engen, J.R., and Guttman, M. (2022). Advances in Hydrogen/Deuterium Exchange Mass Spectrometry and the Pursuit of Challenging Biological Systems. Chem Rev 122, 7562–7623. 10.1021/acs.chemrev.1c00279.

40. Jura, N., Endres, N.F., Engel, K., Deindl, S., Das, R., Lamers, M.H., Wemmer, D.E., Zhang, X., and Kuriyan, J. (2009). Mechanism for activation of the EGF receptor catalytic domain by the juxtamembrane segment. Cell 137, 1293–1307. 10.1016/j.cell.2009.04.025.

41. Zhang, X., Pickin, K.A., Bose, R., Jura, N., Cole, P.A., and Kuriyan, J. (2007). Inhibition of the EGF receptor by binding of MIG6 to an activating kinase domain interface. Nature 450, 741–744. 10.1038/nature05998.

42. Roberts, G. (2015). The role of protein dynamics in allosteric effects-introduction. Biophys Rev 7, 161–163. 10.1007/s12551-015-0174-6.

43. Motlagh, H.N., Wrabl, J.O., Li, J., and Hilser, V.J. (2014). The ensemble nature of allostery. Nature 508, 331–339. 10.1038/nature13001.

44. Schwartz, P.A., Kuzmic, P., Solowiej, J., Bergqvist, S., Bolanos, B., Almaden, C., Nagata, A., Ryan, K., Feng, J., Dalvie, D., et al. (2014). Covalent EGFR inhibitor analysis reveals importance of reversible interactions to potency and mechanisms of drug resistance. Proc Natl Acad Sci U S A 111, 173–178. 10.1073/pnas.1313733111.

45. Fassunke, J., Muller, F., Keul, M., Michels, S., Dammert, M.A., Schmitt, A., Plenker, D., Lategahn, J., Heydt, C., Bragelmann, J., et al. (2018). Overcoming EGFR(G724S)-mediated osimertinib resistance through unique binding characteristics of second-generation EGFR inhibitors. Nat Commun 9, 4655. 10.1038/s41467-018-07078-0.

46. Novotny, C.J., Pollari, S., Park, J.H., Lemmon, M.A., Shen, W., and Shokat, K.M. (2016). Overcoming resistance to HER2 inhibitors through state-specific kinase binding. Nat Chem Biol 12, 923–930. 10.1038/nchembio.2171.

47. Solca, F., Dahl, G., Zoephel, A., Bader, G., Sanderson, M., Klein, C., Kraemer, O., Himmelsbach, F., Haaksma, E., and Adolf, G.R. (2012). Target binding properties and cellular activity of afatinib (BIBW 2992), an irreversible ErbB family blocker. J Pharmacol Exp Ther 343, 342–350. 10.1124/jpet.112.197756.

48. Yan, X.E., Ayaz, P., Zhu, S.J., Zhao, P., Liang, L., Zhang, C.H., Wu, Y.C., Li, J.L., Choi, H.G., Huang, X., et al. (2020). Structural Basis of AZD9291 Selectivity for EGFR T790M. J Med Chem 63, 8502–8511. 10.1021/acs.jmedchem.0c00891.

49. Lamb, Y.N. (2021). Osimertinib: A Review in Previously Untreated, EGFR Mutation-Positive, Advanced NSCLC. Target Oncol 16, 687–695. 10.1007/s11523-021-00839-w.

50. Fu, K., Xie, F., Wang, F., and Fu, L. (2022). Therapeutic strategies for EGFR-mutated non-small cell lung cancer patients with osimertinib resistance. J Hematol Oncol 15, 173. 10.1186/s13045-022-01391-4.

51. Nguyen, K.S., Kobayashi, S., and Costa, D.B. (2009). Acquired resistance to epidermal growth factor receptor tyrosine kinase inhibitors in non-small-cell lung cancers dependent on the epidermal growth factor receptor pathway. Clin Lung Cancer 10, 281–289. 10.3816/CLC.2009.n.039.

52. Cross, D.A., Ashton, S.E., Ghiorghiu, S., Eberlein, C., Nebhan, C.A., Spitzler, P.J., Orme, J.P., Finlay, M.R., Ward, R.A., Mellor, M.J., et al. (2014). AZD9291, an irreversible EGFR TKI, overcomes T790M-mediated resistance to EGFR inhibitors in lung cancer. Cancer Discov 4, 1046–1061. 10.1158/2159-8290.CD-14-0337.

53. Koshland, D.E., Jr. (1998). Conformational changes: how small is big enough? Nat Med 4, 1112–1114. 10.1038/2605.

54. Gutteridge, A., and Thornton, J. (2005). Conformational changes observed in enzyme crystal structures upon substrate binding. J Mol Biol 346, 21–28. 10.1016/j.jmb.2004.11.013.

55. Lopez, E.D., Burastero, O., Arcon, J.P., Defelipe, L.A., Ahn, N.G., Marti, M.A., and Turjanski, A.G. (2020). Kinase Activation by Small Conformational Changes. J Chem Inf Model 60, 821–832. 10.1021/acs.jcim.9b00782.

56. To, C., Beyett, T.S., Jang, J., Feng, W.W., Bahcall, M., Haikala, H.M., Shin, B.H., Heppner, D.E., Rana, J.K., Leeper, B.A., et al. (2022). An allosteric inhibitor against the therapy-resistant mutant forms of EGFR in non-small cell lung cancer. Nat Cancer 3, 402–417. 10.1038/s43018-022-00351-8.

57. Gajiwala, K.S., Feng, J., Ferre, R., Ryan, K., Brodsky, O., Weinrich, S., Kath, J.C., and Stewart, A. (2013). Insights into the aberrant activity of mutant EGFR kinase domain and drug recognition. Structure 21, 209–219. 10.1016/j.str.2012.11.014.

58. Masson, G.R., Burke, J.E., Ahn, N.G., Anand, G.S., Borchers, C., Brier, S., Bou-Assaf, G.M., Engen, J.R., Englander, S.W., Faber, J., et al. (2019). Recommendations for performing, interpreting and reporting hydrogen deuterium exchange mass spectrometry (HDX-MS) experiments. Nat Methods 16, 595–602. 10.1038/s41592-019-0459-y.

59. Houde, D., Berkowitz, S.A., and Engen, J.R. (2011). The utility of hydrogen/deuterium exchange mass spectrometry in biopharmaceutical comparability studies. J Pharm Sci 100, 2071–2086. 10.1002/jps.22432.

60. Agirre, J., Atanasova, M., Bagdonas, H., Ballard, C.B., Basle, A., Beilsten-Edmands, J., Borges, R.J., Brown, D.G., Burgos-Marmol, J.J., Berrisford, J.M., et al. (2023). The CCP4 suite: integrative software for macromolecular crystallography. Acta Crystallogr D Struct Biol 79, 449–461. 10.1107/S2059798323003595.

61. McCoy, A.J., Grosse-Kunstleve, R.W., Adams, P.D., Winn, M.D., Storoni, L.C., and Read, R.J. (2007). Phaser crystallographic software. J Appl Crystallogr 40, 658–674. 10.1107/S0021889807021206.

62. Emsley, P., Lohkamp, B., Scott, W.G., and Cowtan, K. (2010). Features and development of Coot. Acta Crystallogr D Biol Crystallogr 66, 486–501. 10.1107/S0907444910007493.

63. Liebschner, D., Afonine, P.V., Baker, M.L., Bunkoczi, G., Chen, V.B., Croll, T.I., Hintze, B., Hung, L.W., Jain, S., McCoy, A.J., et al. (2019). Macromolecular structure determination using X-rays, neutrons and electrons: recent developments in Phenix. Acta Crystallogr D Struct Biol 75, 861–877. 10.1107/S2059798319011471.

64. Joosten, R.P., Long, F., Murshudov, G.N., and Perrakis, A. (2014). The PDB_REDO server for macromolecular structure model optimization. IUCrJ 1, 213–220. 10.1107/S2052252514009324.

65. Williams, C.J., Headd, J.J., Moriarty, N.W., Prisant, M.G., Videau, L.L., Deis, L.N., Verma, V., Keedy, D.A., Hintze, B.J., Chen, V.B., et al. (2018). MolProbity: More and better reference data for improved all-atom structure validation. Protein Sci 27, 293–315. 10.1002/pro.3330.

