## Supplementary material for "EGFR and tyrosine kinase inhibitor interactions probed by hydrogen-deuterium exchange and mass spectrometry (HDX-MS)": 2024_BioRxiv_YT_Supplemental information.docx

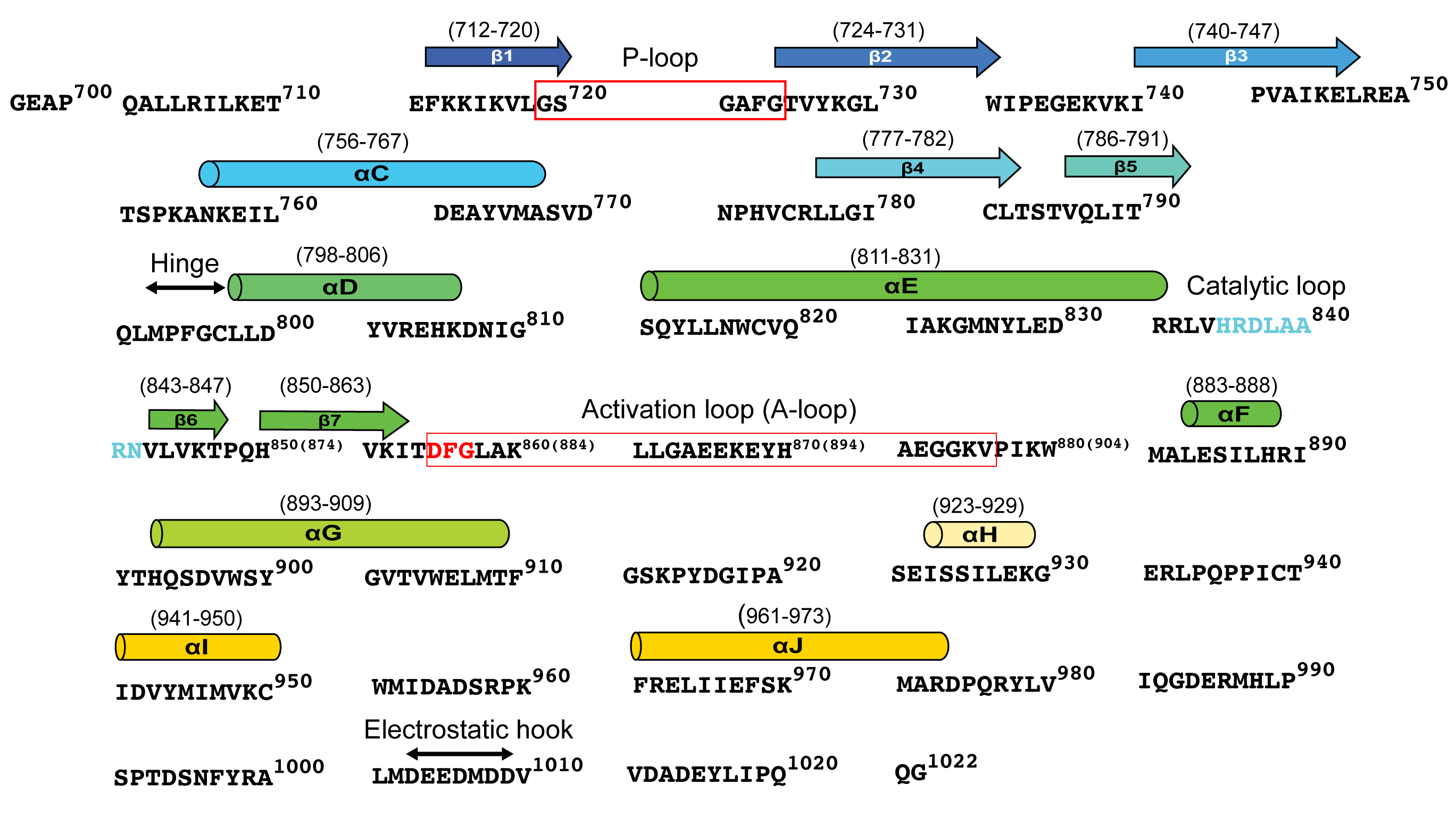

**Figure S1: The amino acid sequence of EGFR tyrosine kinase domain (TKD)**

The amino acid sequence of the TKD with Uniprot EGFR residue number (superscript number above the sequences) is shown. The position of the secondary structures is shown as cartoons above the sequences. The residue number of each secondary structure is shown above the cartoons. The functionally important regions described in the *Results and Discussion* are indicated as shown.

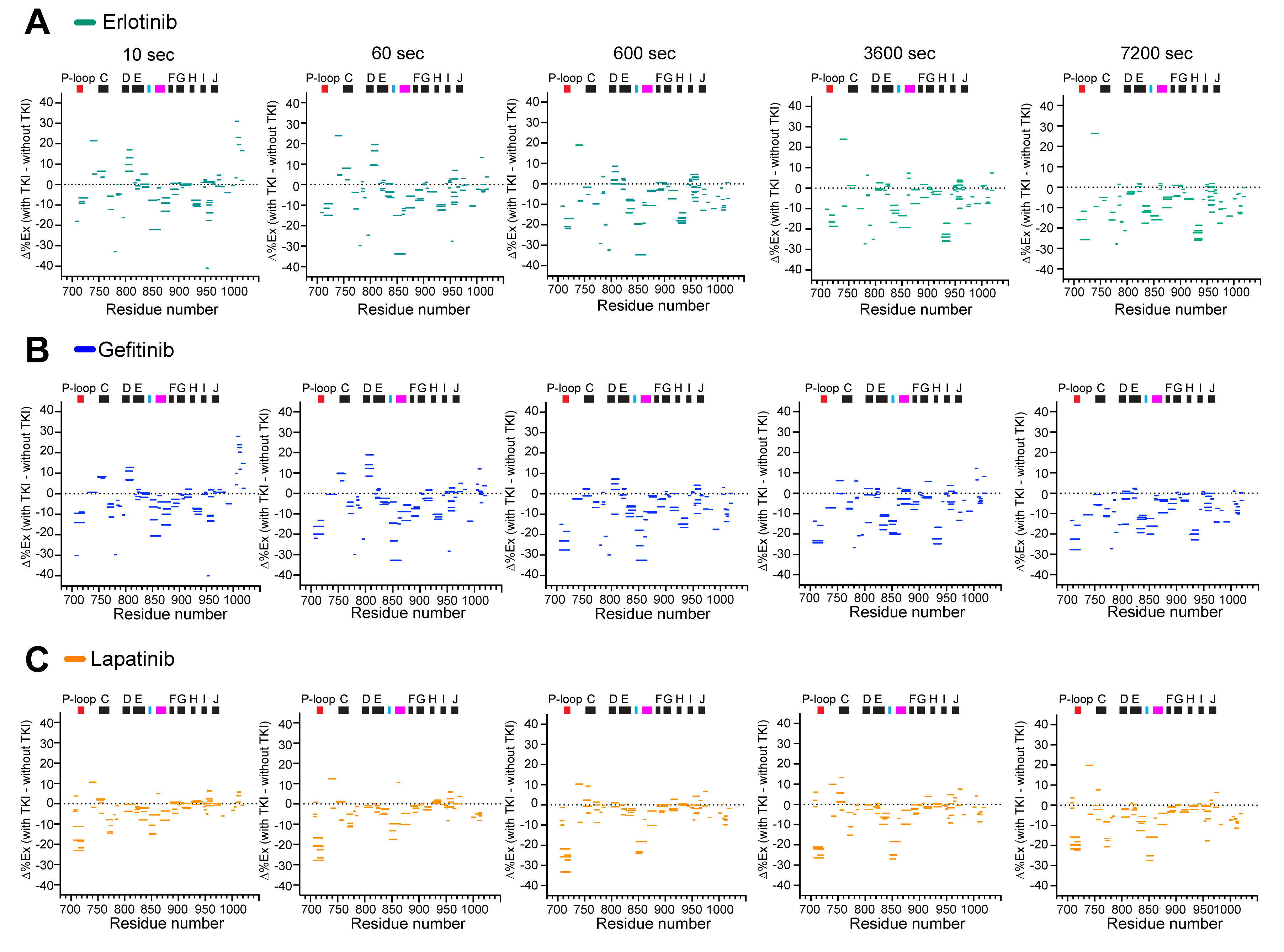

**Figure S2: The dynamics of wild-type EGFR TKD bound to the first-generation TKIs**

Woods plots (Δ%EX map) for the EGFR TKD bound to the first-generation tyrosine kinase inhibitors (TKIs) erlotinib **(A)**, gefitinib **(B)**, or lapatinib **(C)** at the deuterium labeling time of 10, 60, 600, 3600, and 7200 sec at 25^o^C, pD 7.4. The percent exchange differences between the TKI-bound and -unbound wild type TKD (Δ%Ex) of all analyzed peptides, shown as colored horizontal bars, are plotted with the residue number on the X-axis. The regions with positive or negative Δ%Ex values become structurally more flexible or rigid, respectively, with indicated TKIs relative to the TKI-unbound TKD. The position of α-helices (black bar) and functionally important regions, including P-loop, HRD (cyan bar), and DFG·A-loop (magenta bar), are shown at the top of each Δ%EX panel. All HDX data were obtained from n = 3 separate protein preparations, with three independent experiments for each. HDX data statistics are presented in Table S1.

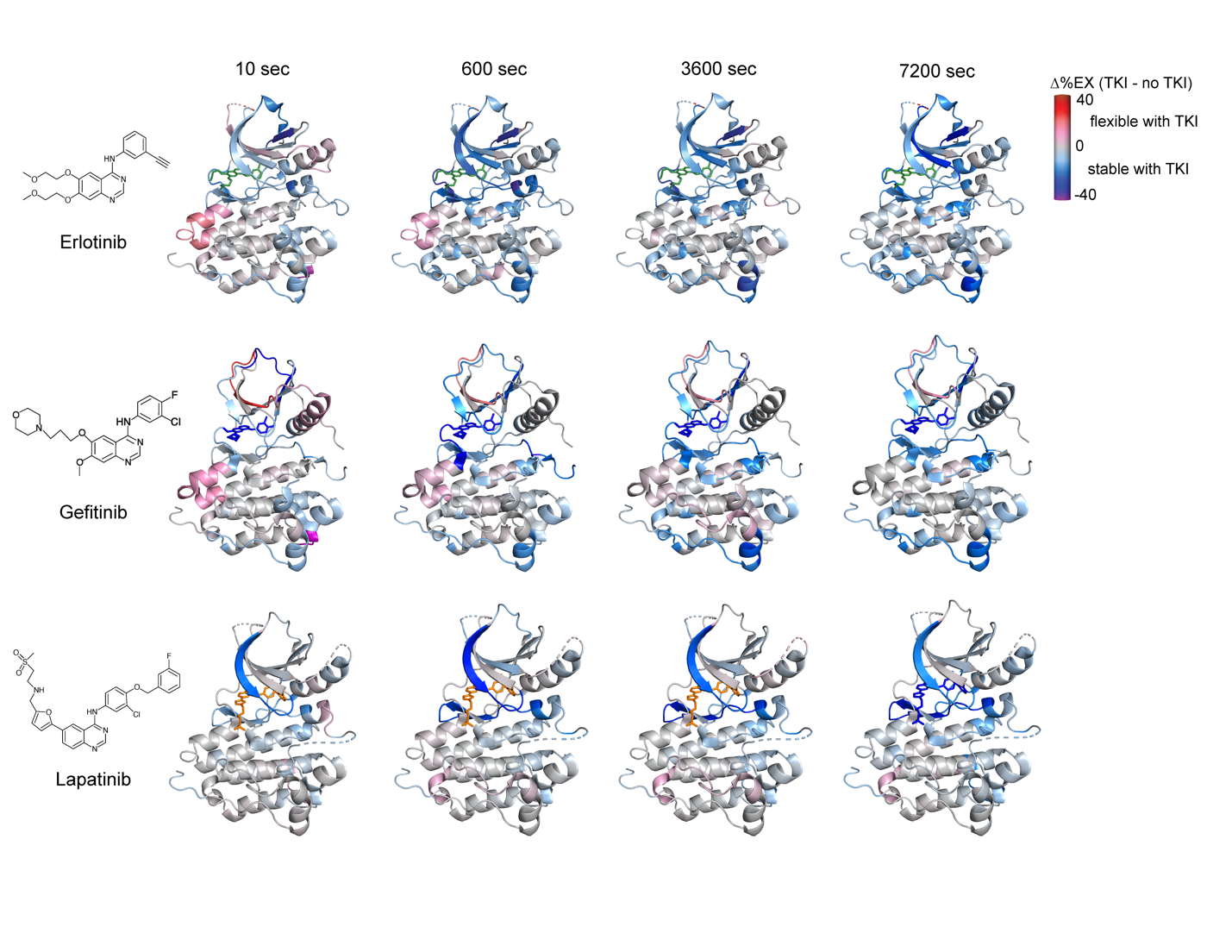

**Figure S3: Structural dynamics of the wild-type TKD with the first-generation TKIs**

The Δ%Ex values of the indicated TKI-bound TKD at the indicated labeling time points are color-coded as shown and mapped onto the EGFR TKD crystal structure bound to erlotinib (PDB ID: 4HJO),^25^ gefitinib (PDB ID:2ITY),^24^ or lapatinib (PDB ID: 1XKK).^6^ The bound TKI is shown as the stick in each figure. All HDX data were obtained from n = 3 separate protein preparations, with three independent experiments for each. The figures are generated using PyMOL (http://www.pymol.org).

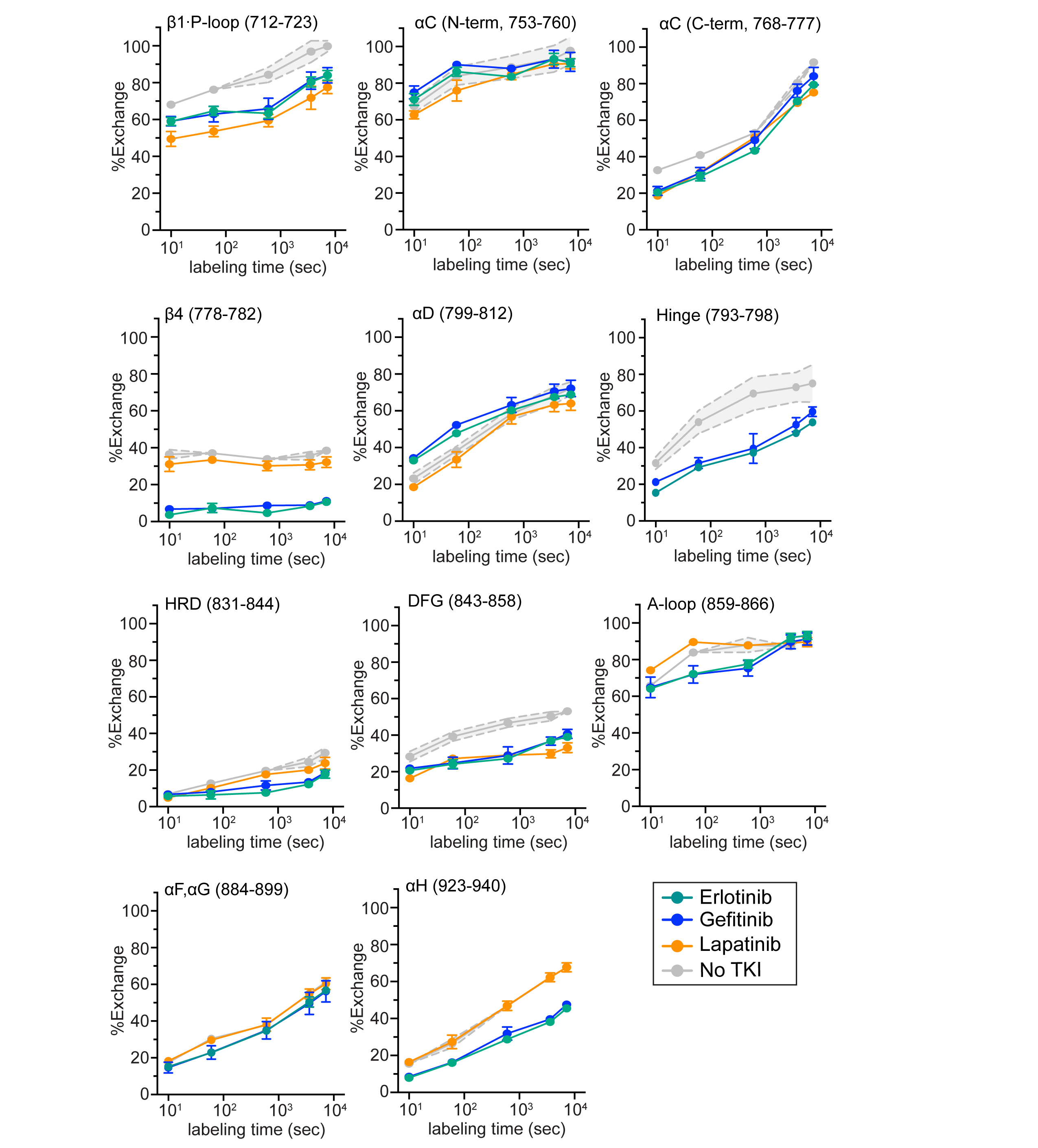

**Figure S4: The percent exchange plots of the indicated regions in the 1st-generation TKI-bound TKD**

The percent exchange of indicated regions for the EGFR TKD without TKI (grey) and with erlotinib (green), gefitinib (blue), or lapatinib (orange) are plotted against the deuterium labeling time (10, 60, 600, 3600, and 7200 sec). The residue number for each region is shown in parentheses. The standard deviations of the unliganded wild-type exchange data are shown as grey error bands. All HDX data were obtained from n = 3 separate protein preparations, with three independent experiments for each.

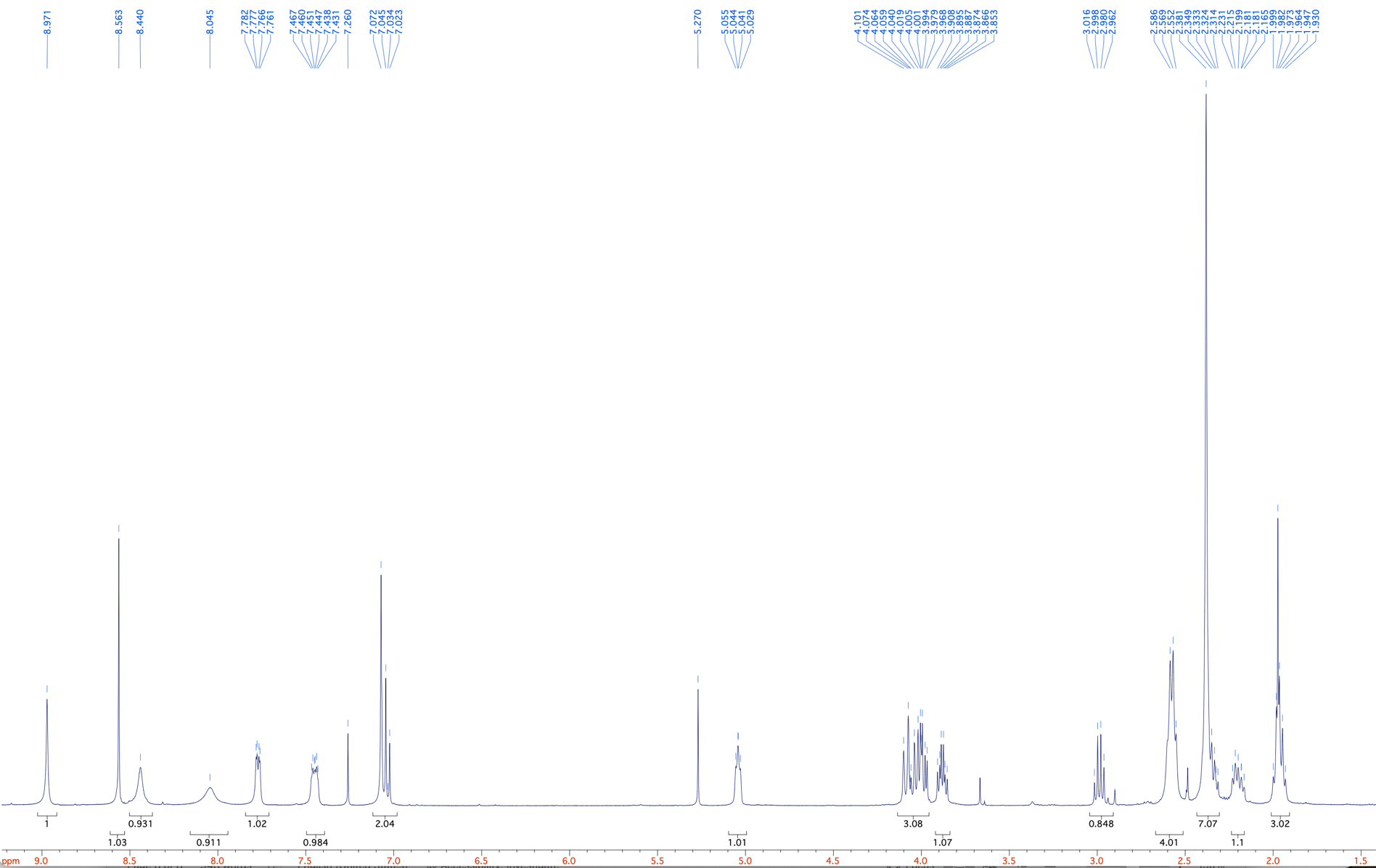

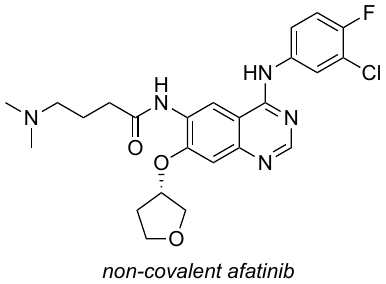

Residual

DCM

**Figure S5: ^1^H NMR spectrum of non-covalent afatinib.** Non-covalent afatinib was synthesized as described in the Methods. The ^1^H peak corresponding to the residual dichloromethane (DCM) is indicated.

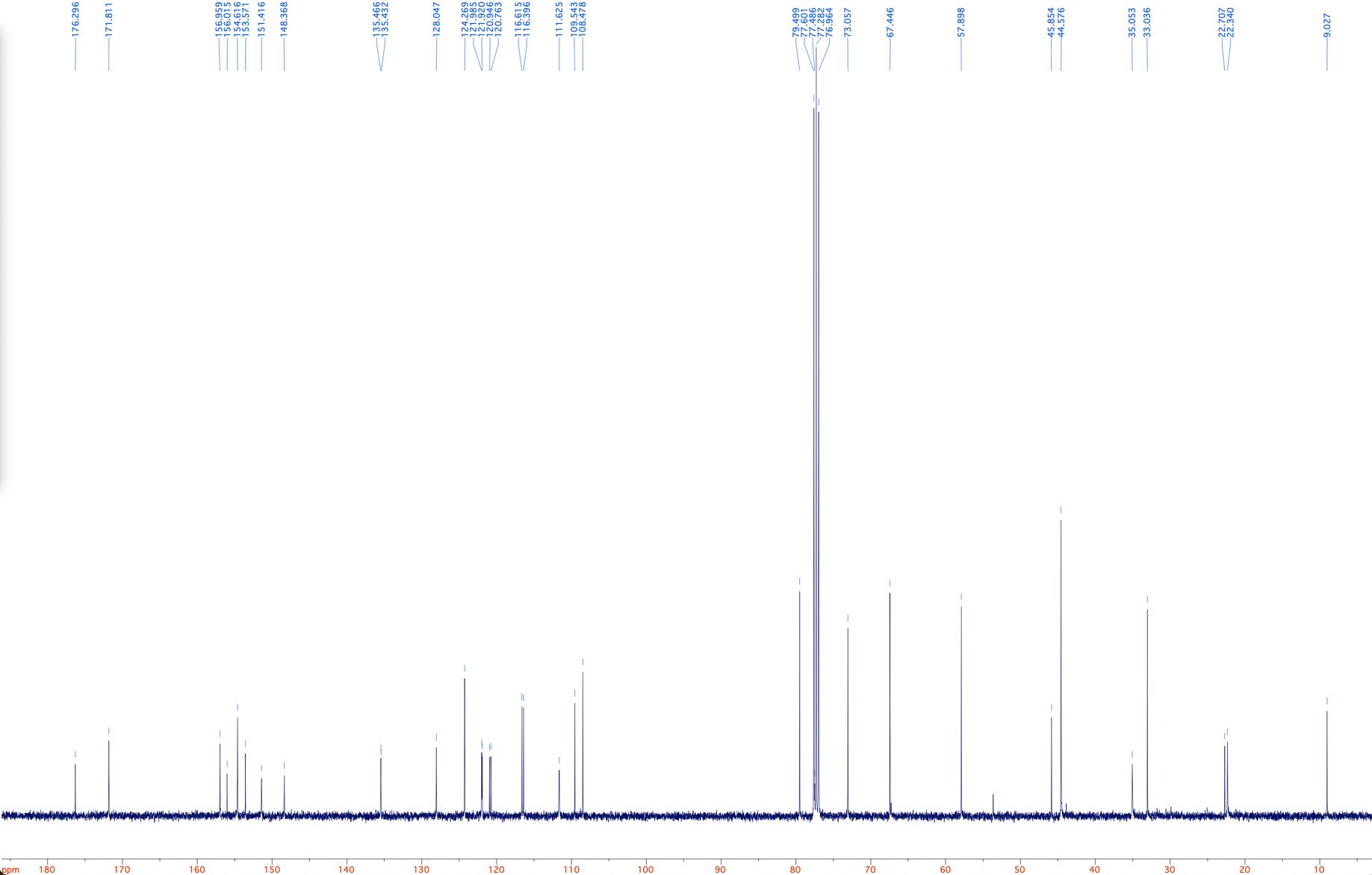

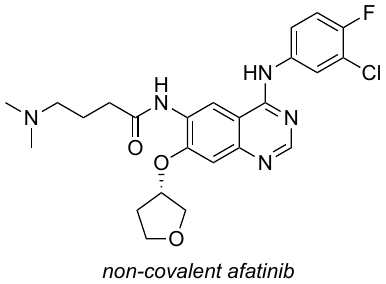

**Figure S6: ^13^C NMR spectrum of non-covalent afatinib.** Non-covalent afatinib was synthesized as described in the Methods.

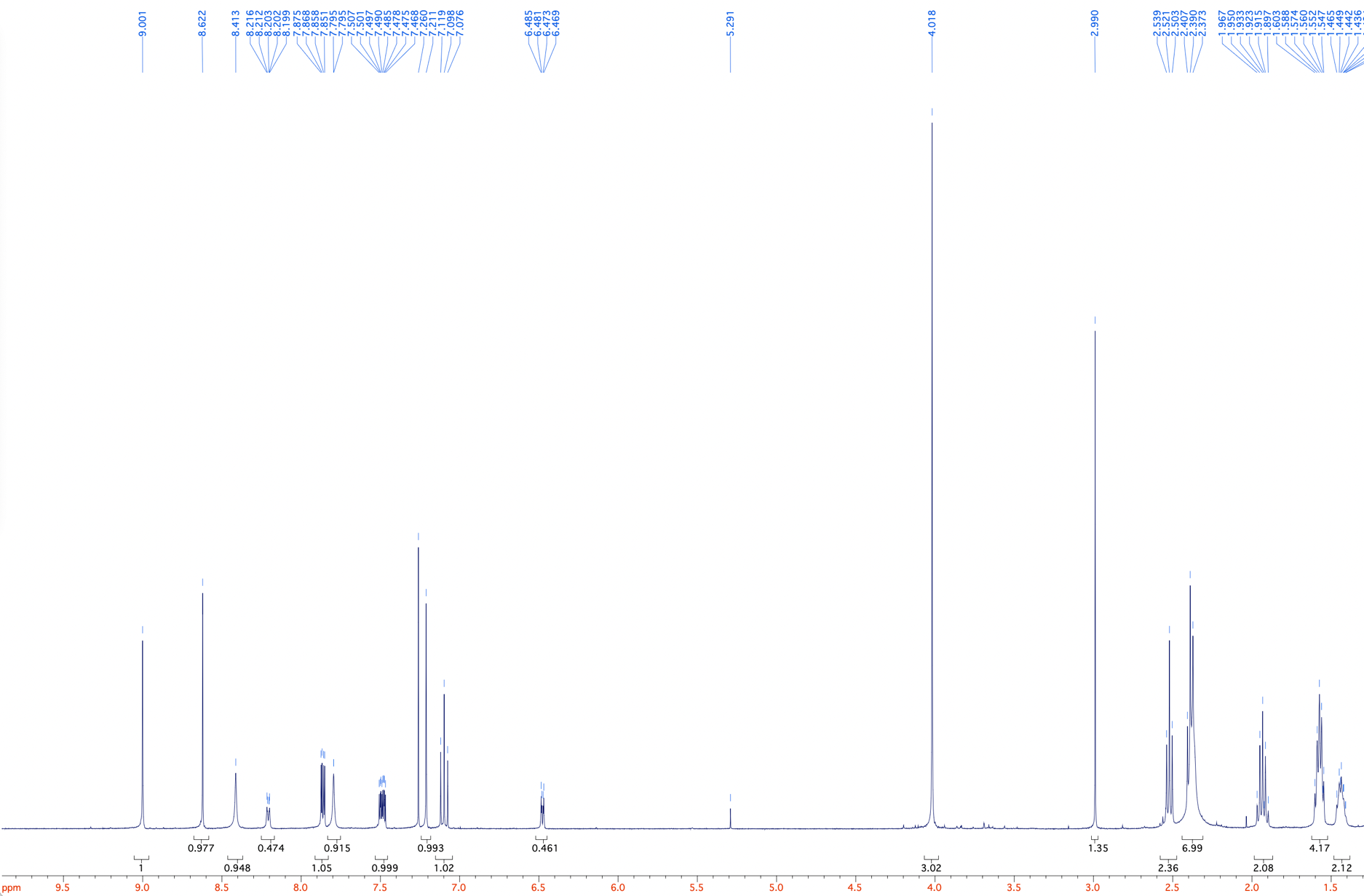

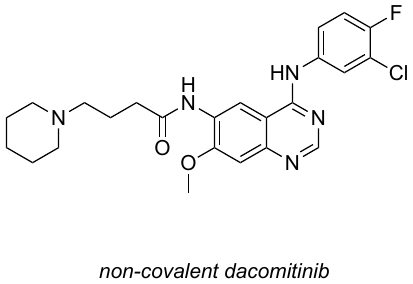

Residual

DCM

Residual

MeOH

**Figure S7: ^1^H NMR spectrum of non-covalent dacomitinib.** Non-covalent dacomitinib was synthesized as described in the Methods. The ^1^H peaks corresponding to the residual dichloromethane (DCM) and methanol are indicated.

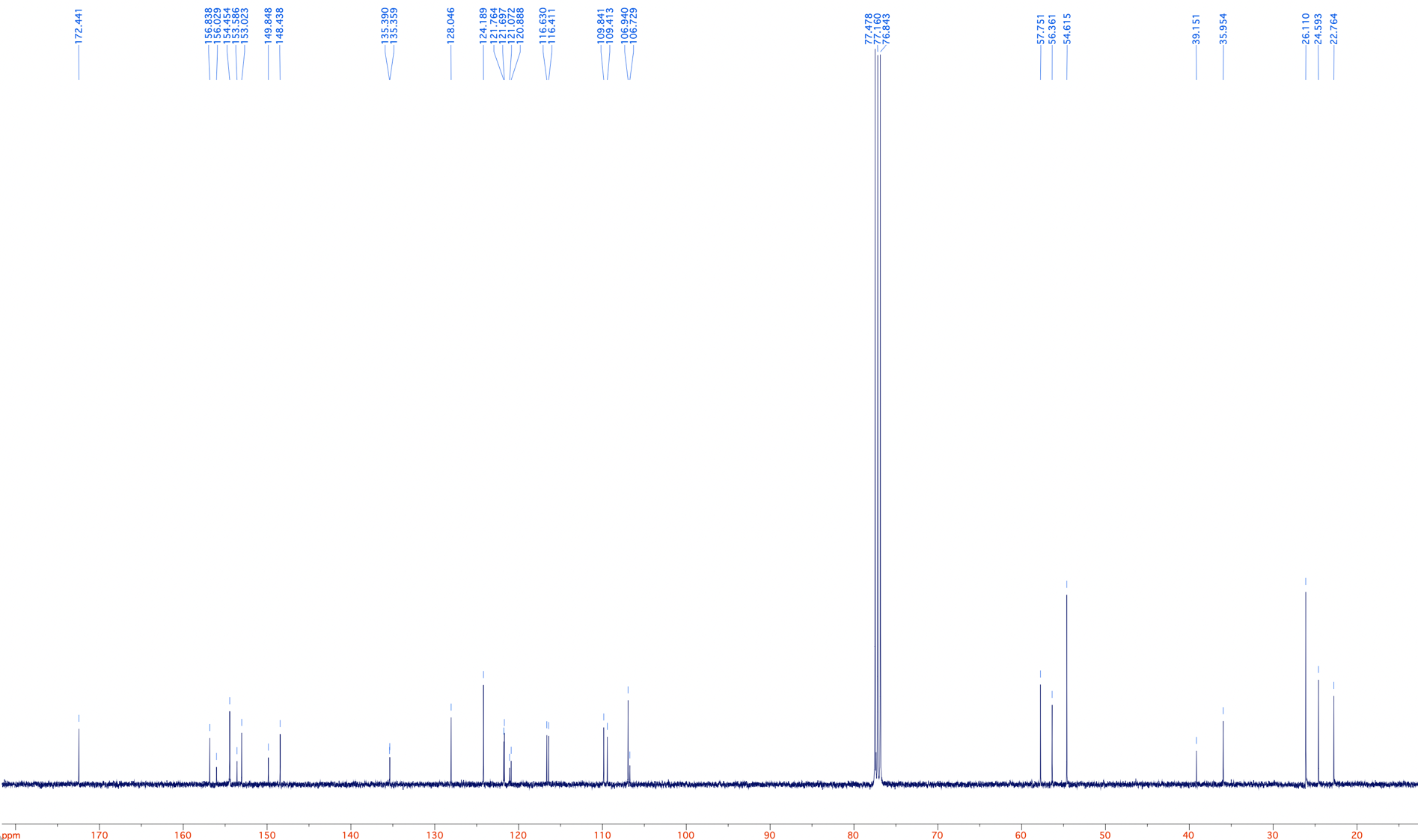

**Figure S8: ^13^C NMR spectrum of non-covalent dacomitinib.** Non-covalent dacomitinib was synthesized as described in the Methods. The ^13^C peaks corresponding to the residual dichloromethane (DCM) and methanol are indicated.

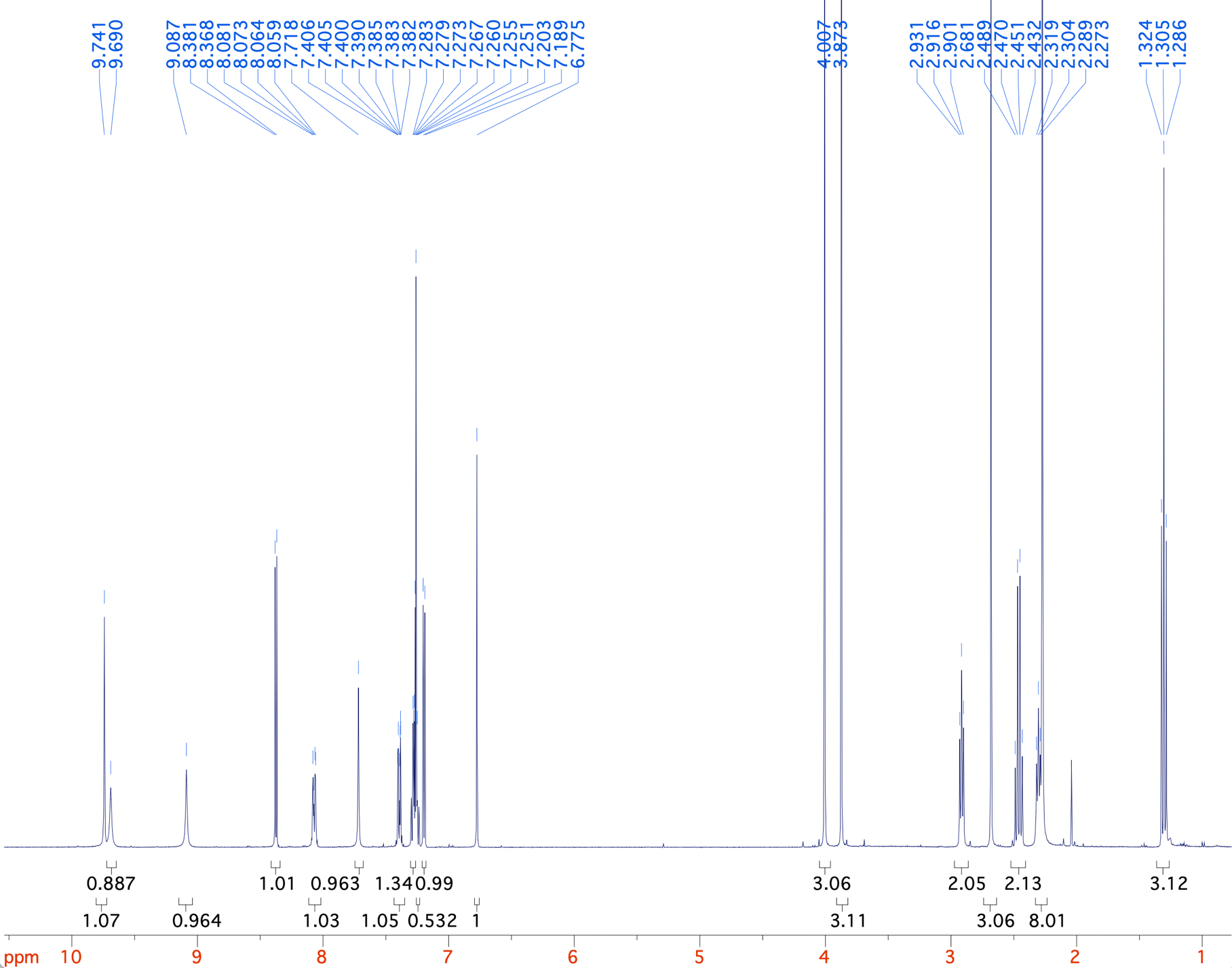

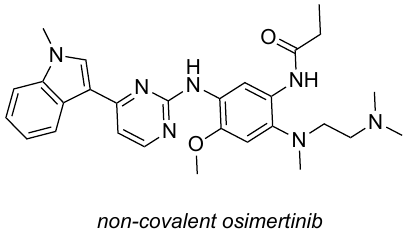

**Figure S9: ^1^H NMR spectrum of non-covalent osimertinib.** Non-covalent osimertinib was synthesized as described in the Methods. The ^1^H peaks corresponding to the residual dichloromethane (DCM) and methanol are indicated.

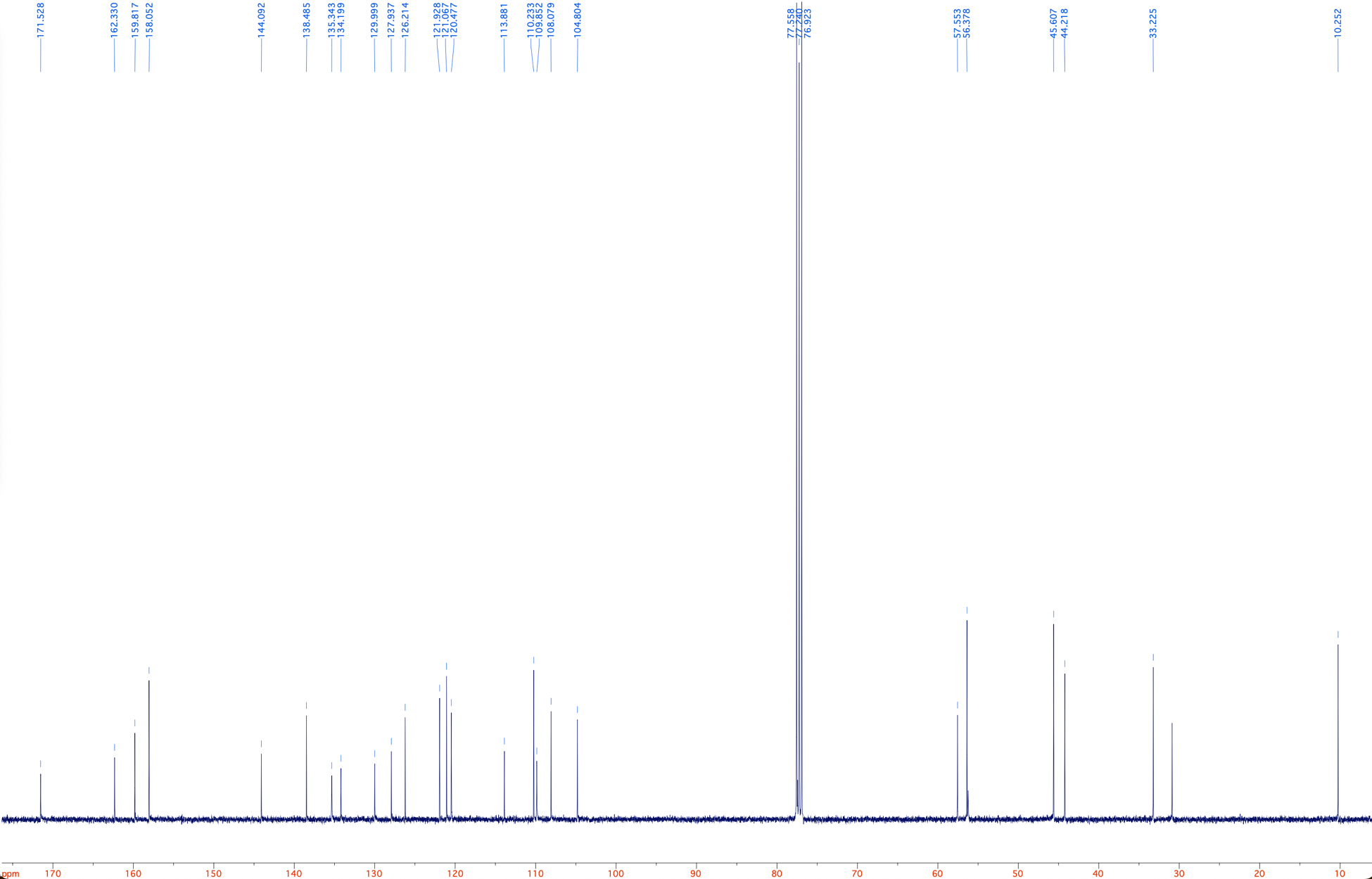

**Figure S10: ^13^C NMR spectrum of non-covalent osimertinib.** Non-covalent osimertinib was synthesized as described in the Methods. The ^13^C peaks corresponding to the residual dichloromethane (DCM) and methanol are indicated.

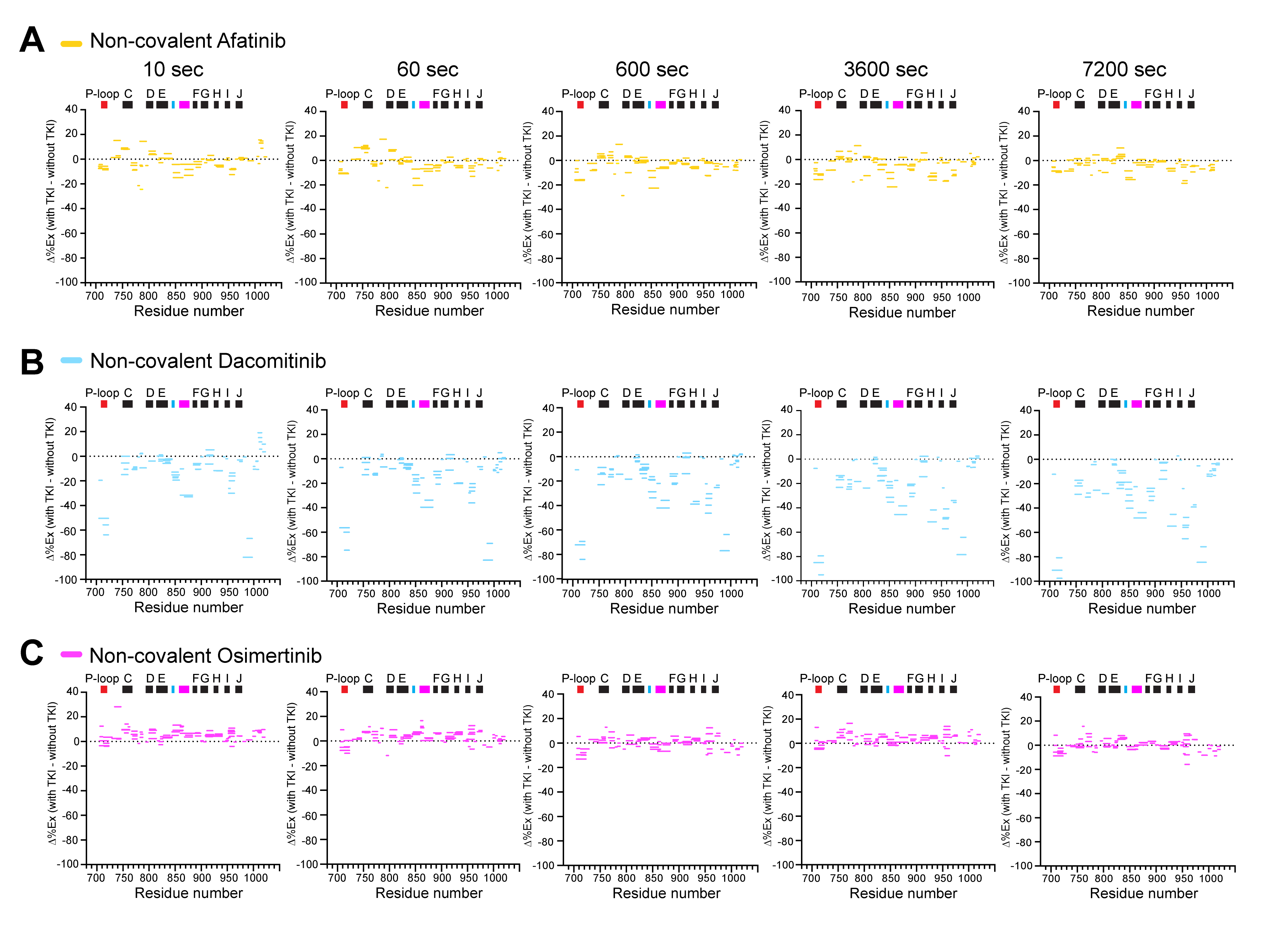

**Figure S11: The dynamics of wild-type EGFR TKD bound to non-covalent second- and third-generation TKIs**

Woods plots (Δ%EX map) for the EGFR TKD bound to non-covalent afatinib **(A)**, non-covalent dacomitinib **(B)**, or non-covalent osimertinib **(C)** at the deuterium labeling time of 10, 60, 600, 3600, and 7200 sec at 25^o^C, pD 7.4. The percent exchange differences between the TKI-bound and -unbound wild type TKD (Δ%Ex) of all analyzed peptides, shown as colored horizontal bars, are plotted with the residue number on the X-axis. The regions with positive or negative Δ%Ex values become structurally more flexible or rigid, respectively, with indicated TKIs relative to the TKI-unbound TKD. The position of α-helices (black bar) and functionally important regions, including P-loop, HRD (cyan bar), and DFG·A-loop (magenta bar), are shown at the top of each Δ%EX map. All HDX data were obtained from n = 3 separate protein preparations except for the non-covalent dacomitinib-TKD (n =2 separate protein preparations), with three independent experiments for each. HDX data statistics are presented in Table S2.

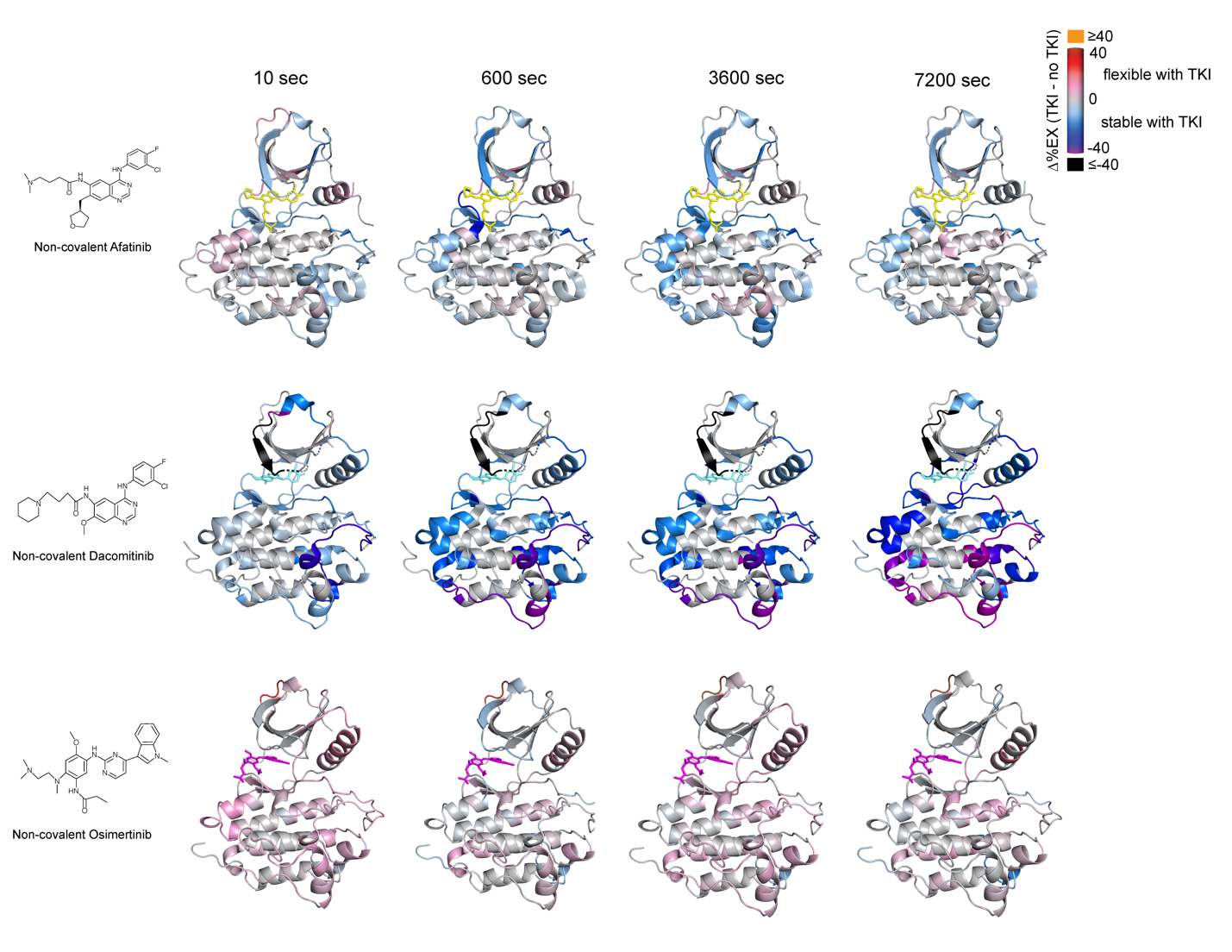

**Figure S12:** **The dynamics of the wild-type EGFR TKD encounter complex**

The Δ%EX values of peptides in each encounter complex are color-coded and mapped onto crystal structures of wild-type EGFR TKD with non-covalent afatinib (PDB ID: 4G5J),^47^ non-covalent dacomitinib (PDB ID:4I23),^57^ or non-covalent osimertinib (from this study). The bound inhibitor is shown as sticks in each figure. The figures are generated using PyMOL (http://www.pymol.org).

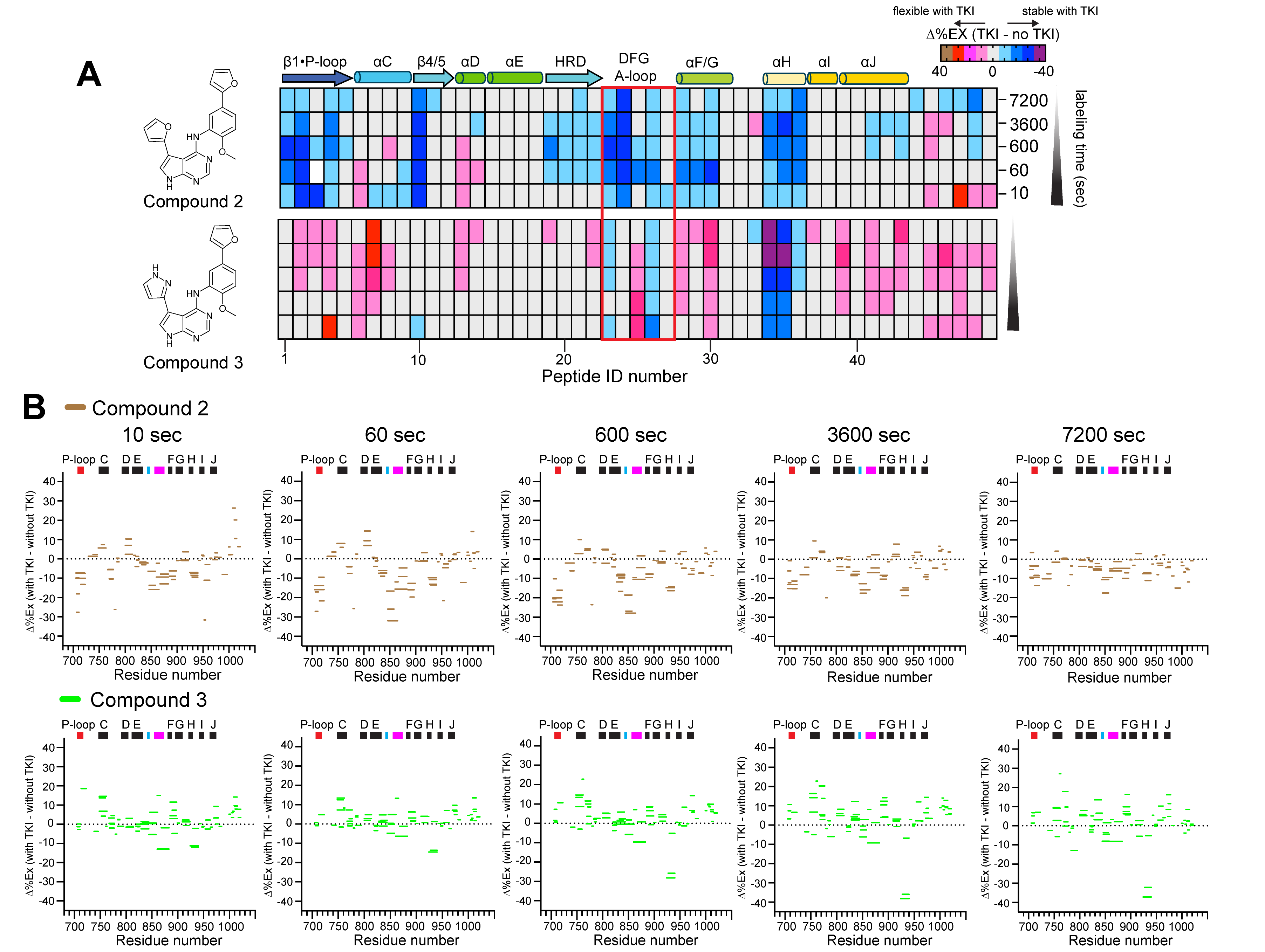

**Figure S13: The overall dynamics of compound 2- or 3-bound EGFR TKD**

**(A)** The Δ%EX values of the same peptides derived from compound 2- or 3-bound TKD are color-coded as shown on the right and mapped from the N-terminal peptide to the C-terminal peptide (#1 to #49). The approximate position of the secondary structures is shown as cartoons at the top of the heat map. The Δ%EX values are shown from the bottom (10 sec) to the top (7200 sec) of each heat map. *Supplemental information* lists the amino acid sequence and Δ%EX value of each peptide ID. **(B)** Woods plots (Δ%EX map) of the compound 2- or 3-bound TKD at the indicated deuterium labeling time. The percent exchange differences between the TKI-bound and -unbound wild type TKD (Δ%Ex) of each peptide, shown as colored horizontal bars, are plotted with the residue number on the X-axis. The position of α-helices (black bar) and functionally important regions, including P-loop, HRD (cyan bar), and DFG·A-loop (magenta bar), are shown at the top of each Δ%EX map. All HDX data were obtained from n = 3 (compound 2) or n = 2 (compound 3) separate protein preparations, with three independent experiments for each. HDX data statistics are presented in Table S3. The structure maps are generated using PyMOL (http://www.pymol.org).

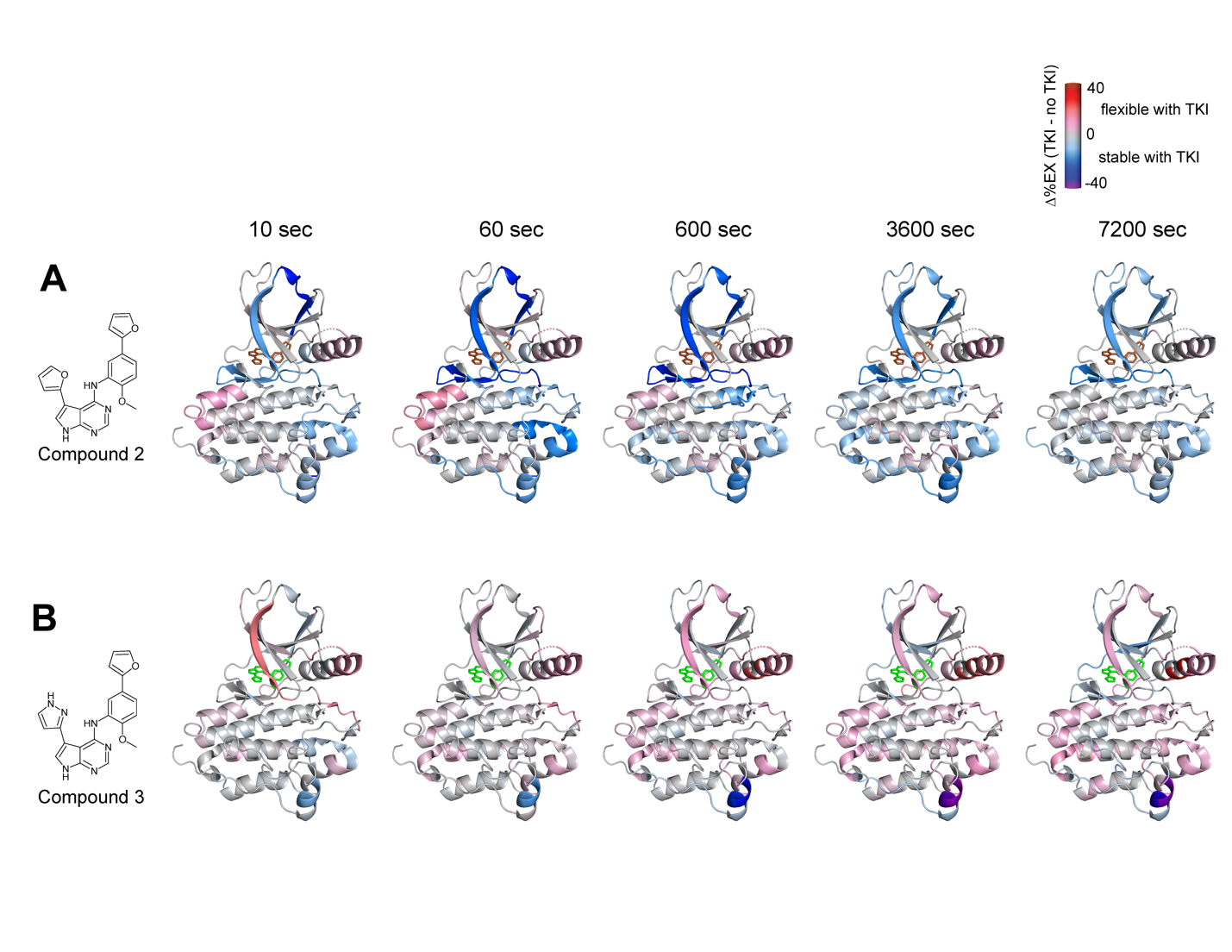

**Figure S14: Structural dynamics of the EGFR TKD bound to compound 2 or 3**

The Δ%Ex values of peptides in each TKI-bound TKD are color-coded and mapped onto a crystal structure of wild-type EGFR TKD with compound 2 **(A)** (PDB ID: 5JEB).^46^ The bound compound 2 is shown as brown sticks. Because a crystal structure of the EGFR TKD with compound 3 is unavailable, the compound 2-bound TKD structure was used to map the %EX values of the compound 3-bound TKD **(B)**. The figures are generated using PyMOL (http://www.pymol.org).

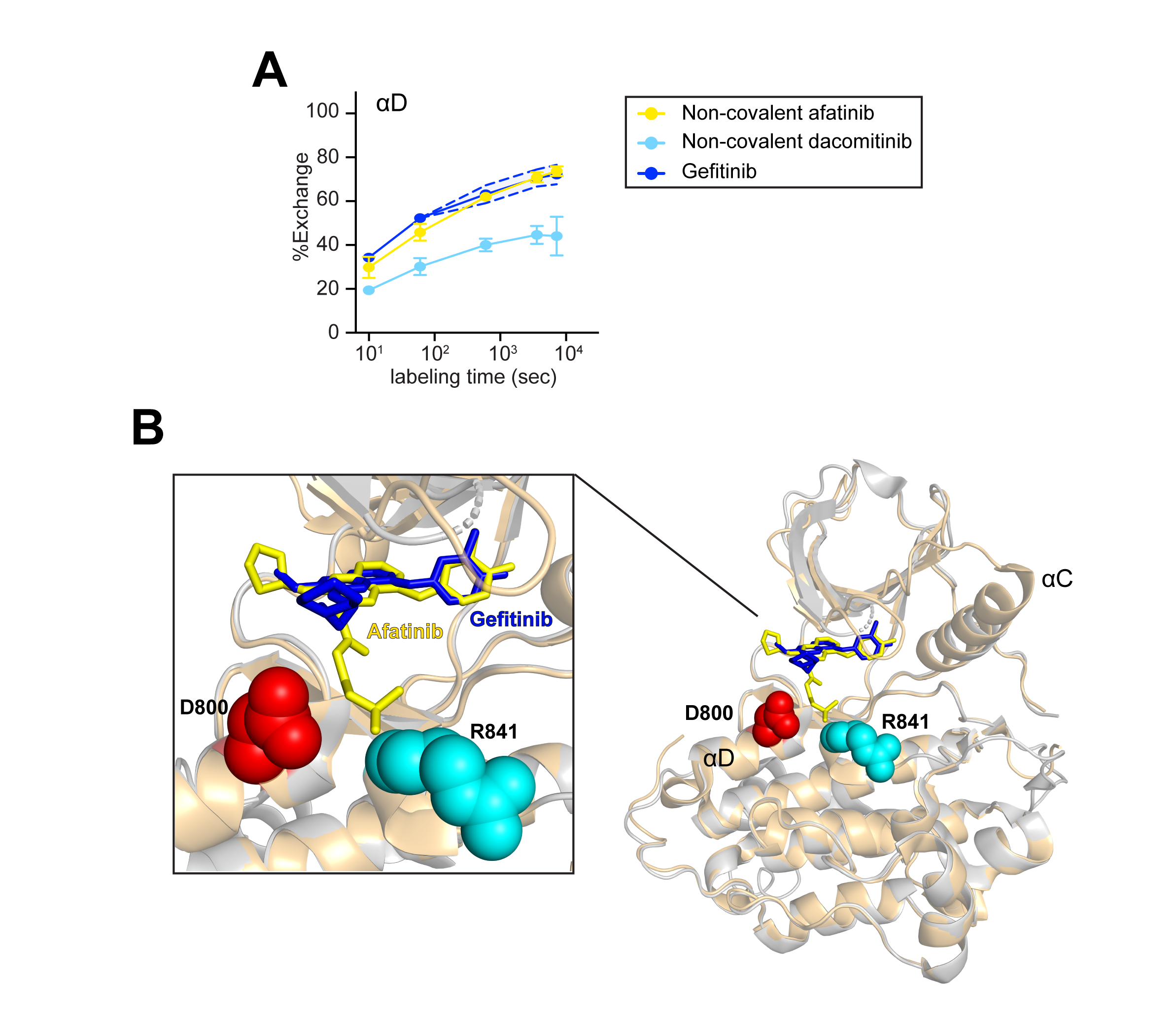

**Figure S15: Structural fluctuations of αD propagate to the β4-β5 hydrophobic pocket via a bound TKI**

**(A)** The percent exchange of αD (residue 799-812) in the wild-type TKD bound to non-covalent afatinib (yellow), non-covalent dacomitinib (light blue), or gefitinib (blue) is plotted against the deuterium labeling time (10, 60, 600, 3600, and 7200 sec). The standard deviation of the percent exchange of the gefitinib-bound TKD is shown as dotted error bands. **(B)** Superposed crystal structures of the wild-type TKD bound to afatinib (PDB ID: 4G5J)^47^ and gefitinib (PDB ID: 2 ITY)^24^ are shown as grey and orange cartoons, respectively. The side chains of D800 and R841 are shown as red and cyan spheres, respectively. The bound afatinib (yellow) and gefitinib (blue) are shown as sticks.

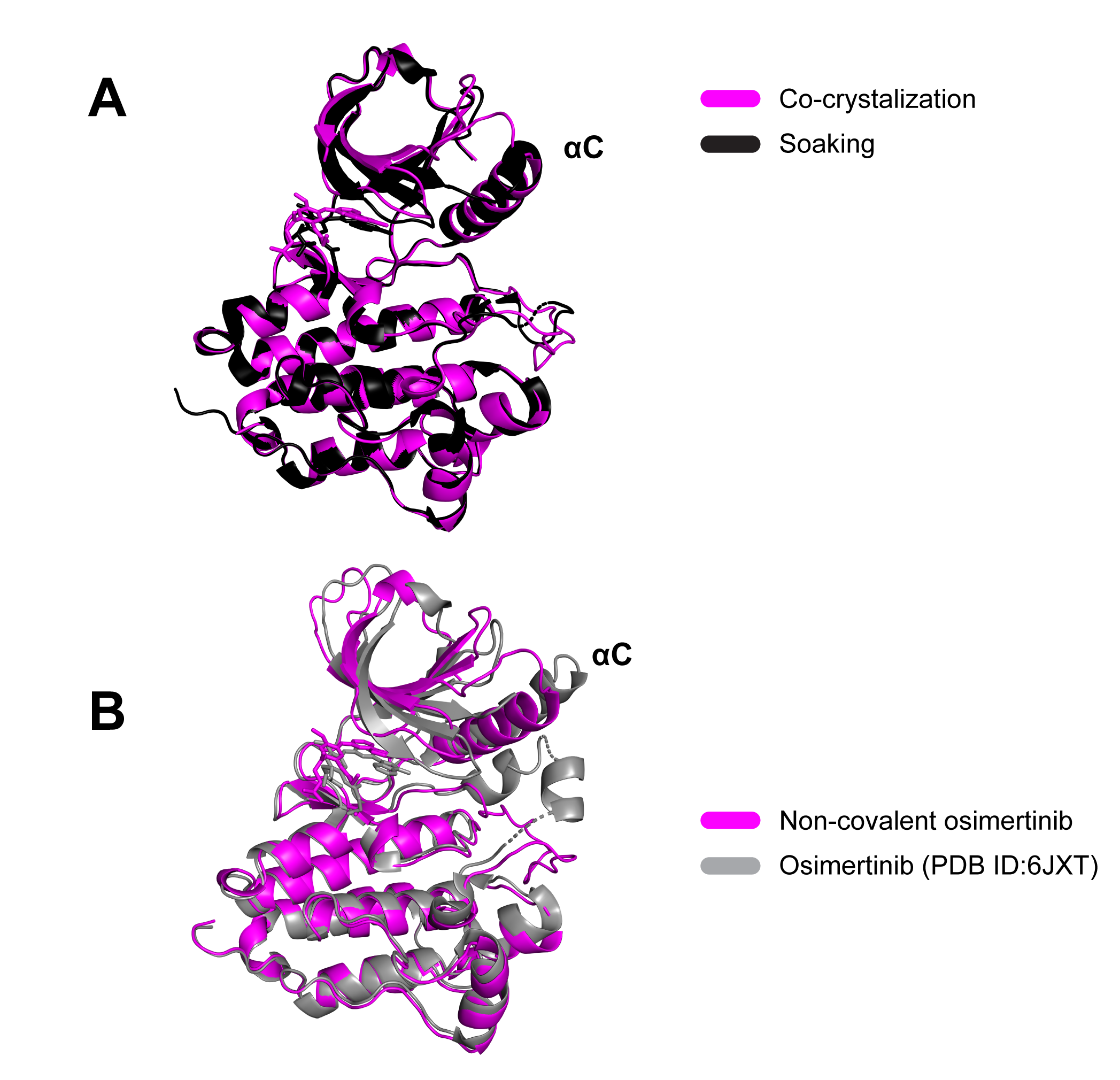

**Figure S16: The encounter to covalent complex formation involves structural rearrangement of N-lobe β-strands and A-loop**

**(A)** Superposed crystal structures of wild-type EGFR TKD complexed with non-covalent osimertinib solved by co-crystallization (magenta cartoon, PDB ID: XXX) or soaking (black cartoon, PDB ID: XXX) **(B)** Superposed crystal structures of wild-type EGFR TKD bound to non-covalent osimertinib (PDB ID:XXXX) solved by co-crystallization and covalent osimertinib (PDB ID: 6JXT)^48^ are shown as magenta and black cartoons, respectively. The bound inhibitor is shown as sticks. In the structure of the wild-type encounter complex, disordered N- and C-terminal regions (residue 695-703 and 981-1010) are not shown for clarity.

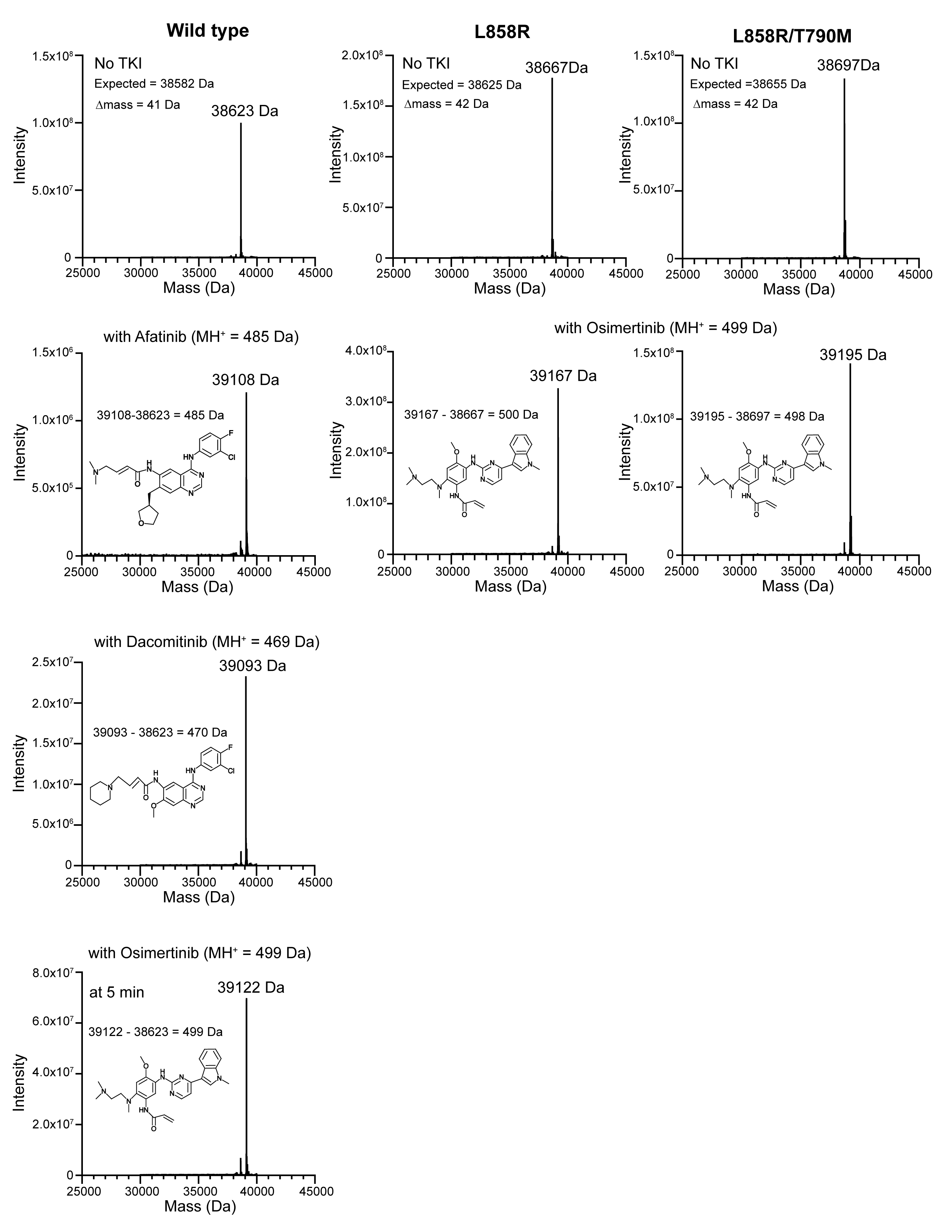

**Figure S17: Intact mass analysis of wild type, L858R, and L858R/T790M covalent complexes**

The intact mass analysis of the wild-type, L858R, and L858R/T790M by mass spectrometry. The MS spectra of purified proteins were acquired without and with indicated inhibitors after 5 min (wild type) or 1 hour (L858R and L858R/T790M) TKI incubation at 25^o^C in 20 mM HEPES, 100 mM NaCl, pH 7.4 as described in Methods. All MS spectra were deconvoluted using MassLynx MaxEnt1 (Waters). The mass difference of 41-42 Dalton of each protein without TKI is attributed to the N-acetylation of purified proteins.

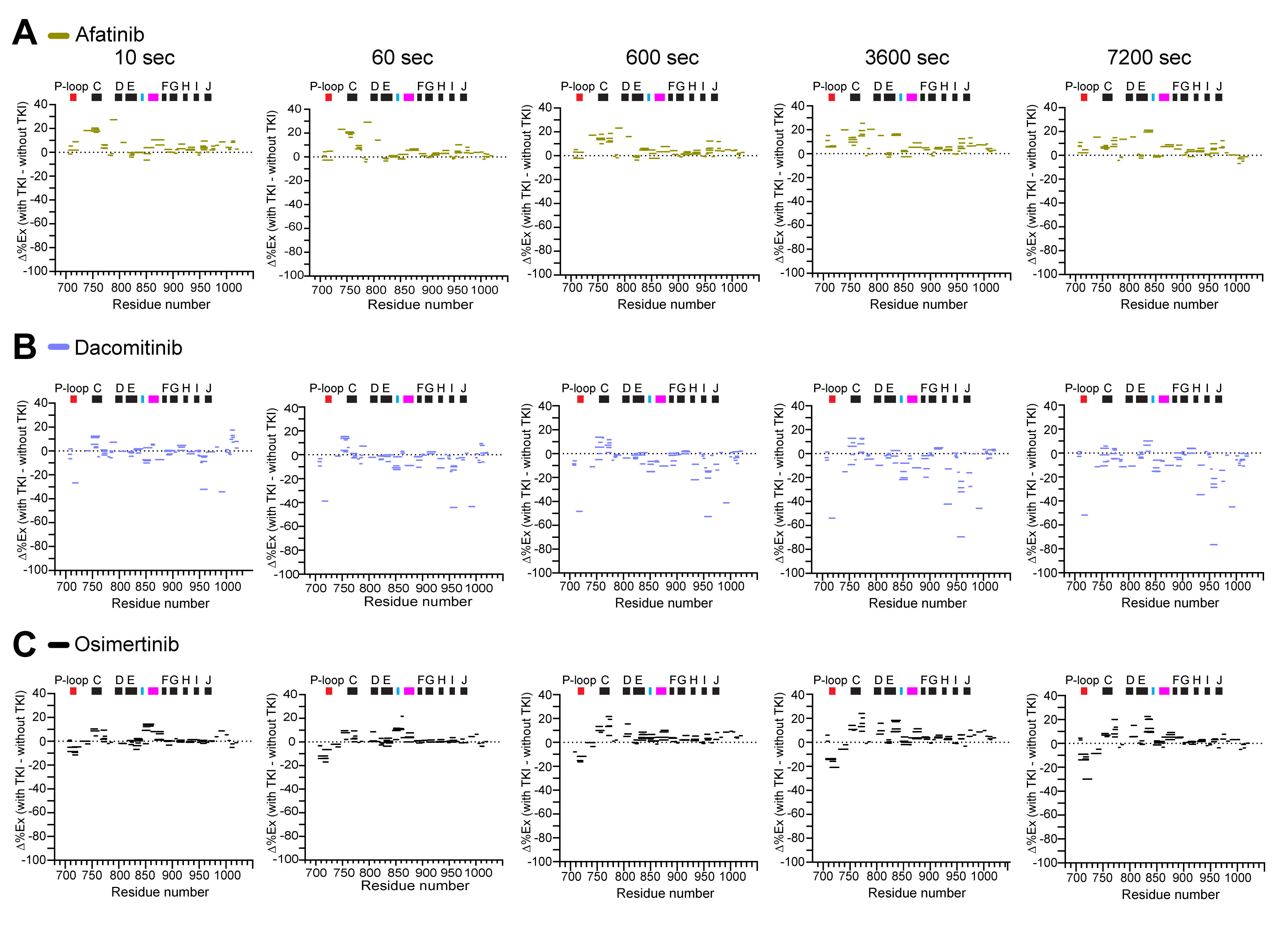

**Figure S18: Woods Plots of covalent TKI-EGFR TKD complexes**

Woods plots (Δ%EX map) for the EGFR TKD complexed with covalent afatinib **(A)**, dacomitinib **(B)**, or osimertinib **(C)** at the deuterium labeling time of 10, 60, 600, 3600, and 7200 sec at 25^o^C, pD 7.4. The percent exchange differences between the TKI-bound and -unbound wild type TKD (Δ%Ex) of all analyzed peptides, shown as colored horizontal bars, are plotted with the residue number on the X-axis. The regions with positive or negative Δ%Ex values become structurally more flexible or rigid, respectively, with indicated TKIs relative to the TKI-unbound TKD. The position of α-helices (black bar) and functionally important regions, including P-loop, HRD (cyan bar), and DFG·A-loop (magenta bar), are shown at the top of each Δ%EX map. All HDX data were obtained from n = 3 separate protein preparations except for dacomitinib-TKD (n = 2 separate protein preparations), with three independent experiments for each. HDX data statistics are presented in Table S5.

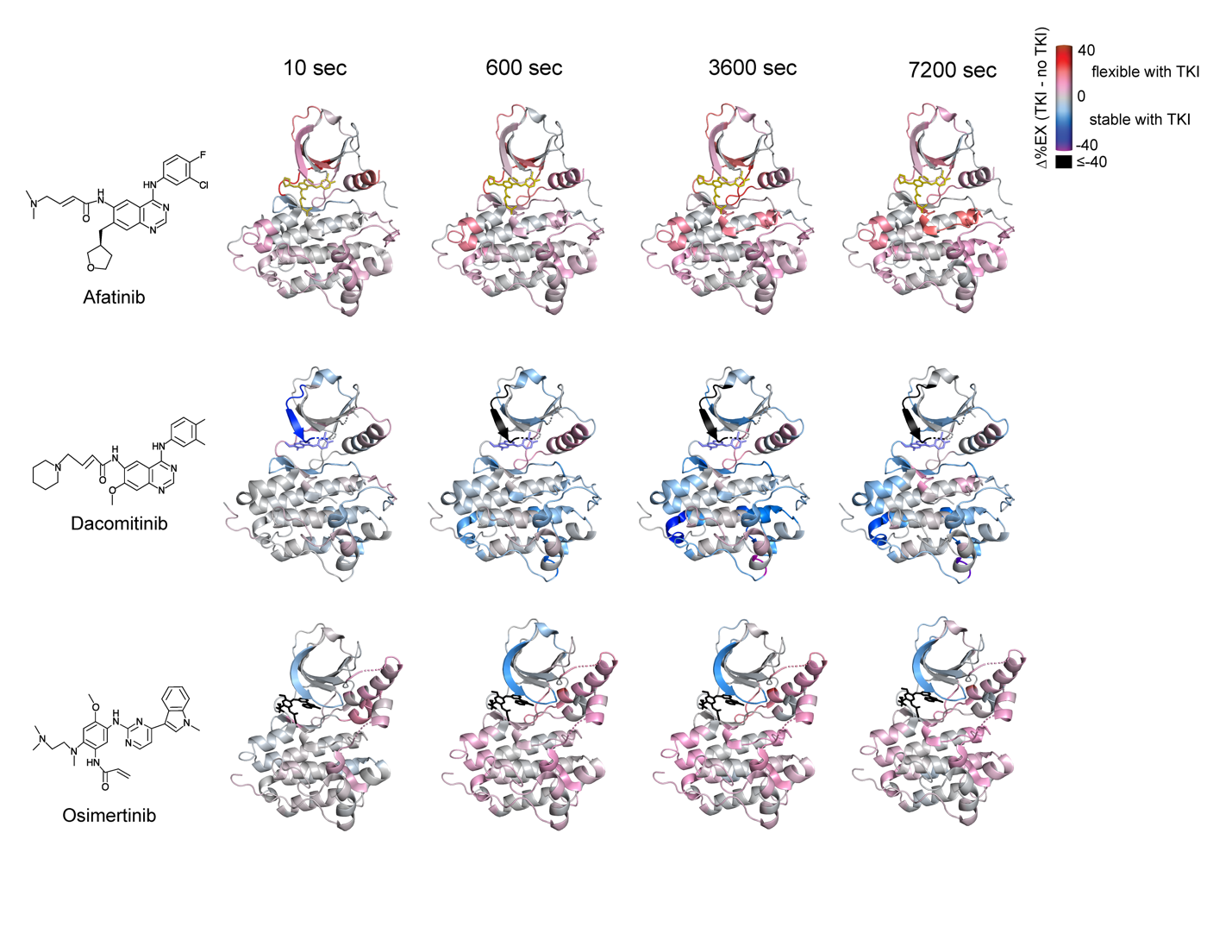
**Figure S19:** **Structural dynamics of the EGFR TKD complexed with 2nd- or 3rd-generation covalent inhibitor**

The Δ%Ex values are color-coded and mapped onto crystal structures of wild-type EGFR TKD afatinib (PDB ID: 4G5J),^47^ dacomitinib (PDB ID: 4I23),^57^ or osimertinib (PDB ID: 6JXT)^48^ covalent complex at the indicated deuterium labeling time. The bound inhibitor is indicated as a stick in each figure. The figures are generated using PyMOL (http://www.pymol.org).

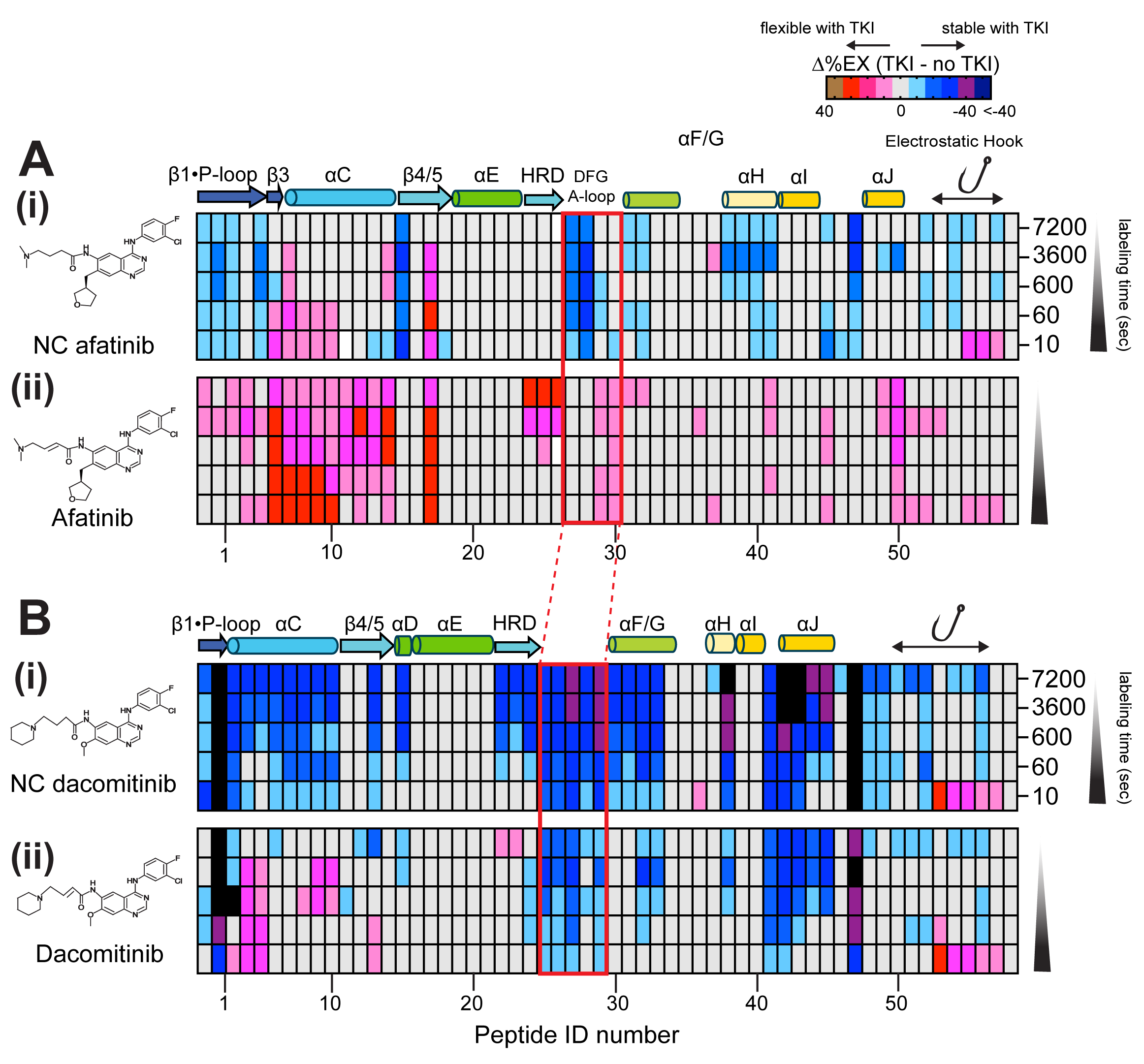

**Figure S20: Dynamic transition from the encounter to covalent complexes**

**(A)** The Δ%EX heatmaps (TKI - no TKI) of the wild-type TKD complexed with non-covalent afatinib **(i)** and covalent afatinib **(ii)**, **(B)** non-covalent dacomitinib (i) and covalent dacomitinib (ii) are shown. The columns show color-coded Δ%EX of the same peptides in the inhibitor complexes at the indicated labeling times. Peptides are assigned peptide ID numbers from the N-terminus (left) to C-terminus (right) on the X-axis. *Supplemental information* lists the amino acid sequence and Δ%EX value of each peptide ID. The position of the secondary structures and conserved HRD and DFG motif regions are shown at the top of the heatmap. The red square box on the heatmaps indicates peptides containing the DFG motif and A-loop.

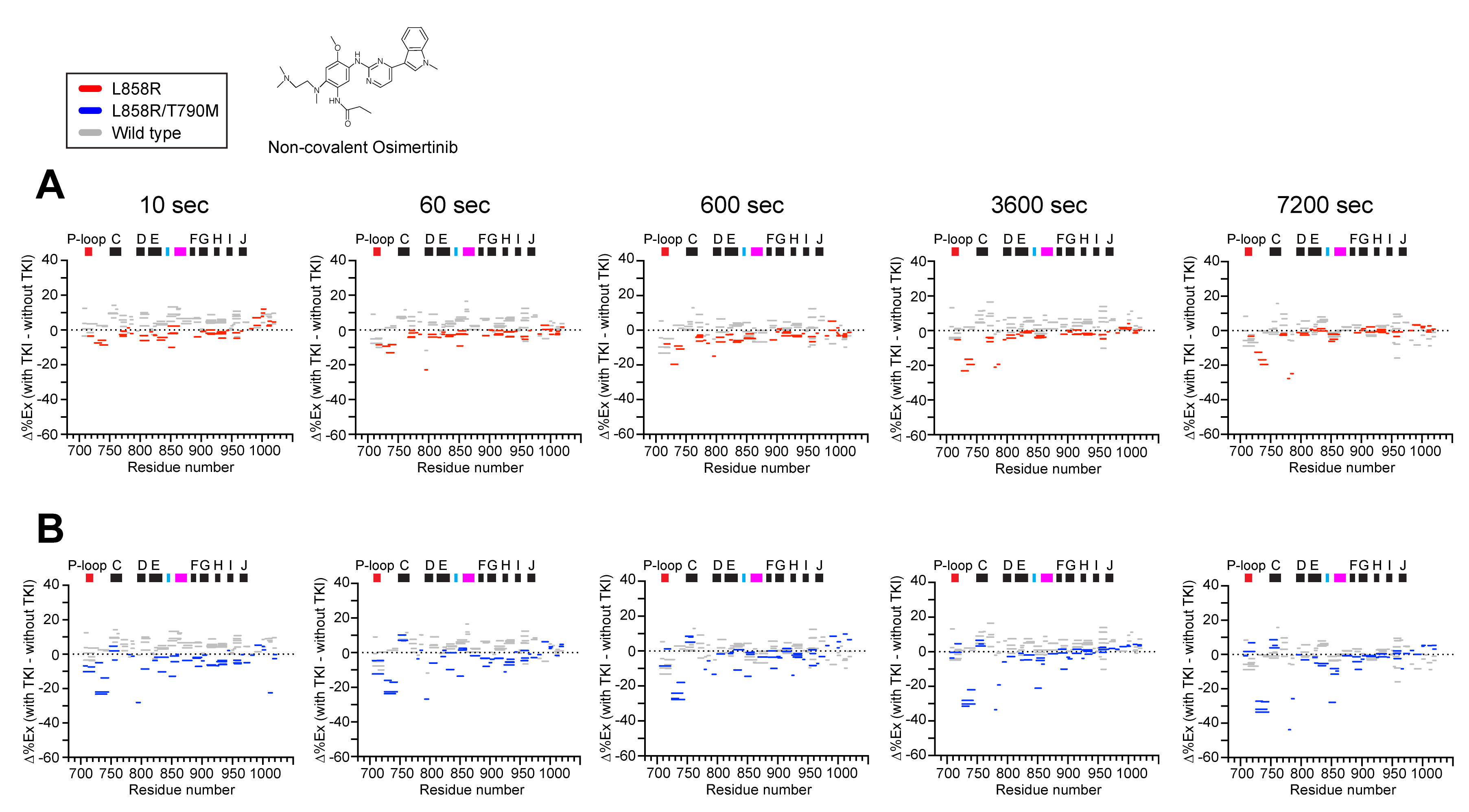

**Figure S21: Woods Plots of non-covalent osimertinib-EGFR TKD complexes**

Woods plots (Δ%EX map) for EGFR TKD bound to non-covalent osimertinib at the deuterium labeling time of 10, 60, 600, 3600, and 7200 sec at 25^o^C, pD 7.4. The percent exchange differences between the TKI-bound and -unbound TKD (Δ%Ex) of all analyzed peptides, shown as colored horizontal bars, are plotted with the residue number on the X-axis. The regions with positive or negative Δ%Ex values become structurally more flexible or rigid, respectively, with indicated TKIs relative to the TKI-unbound TKD. The position of α-helices (black bar) and functionally important regions, including P-loop, HRD (cyan bar), and DFG·A-loop (magenta bar), are shown at the top of each Δ%EX map. All HDX data were obtained from n = 3 separate protein preparations, with three independent experiments for each. HDX data statistics are presented in Tables S6 and S7.

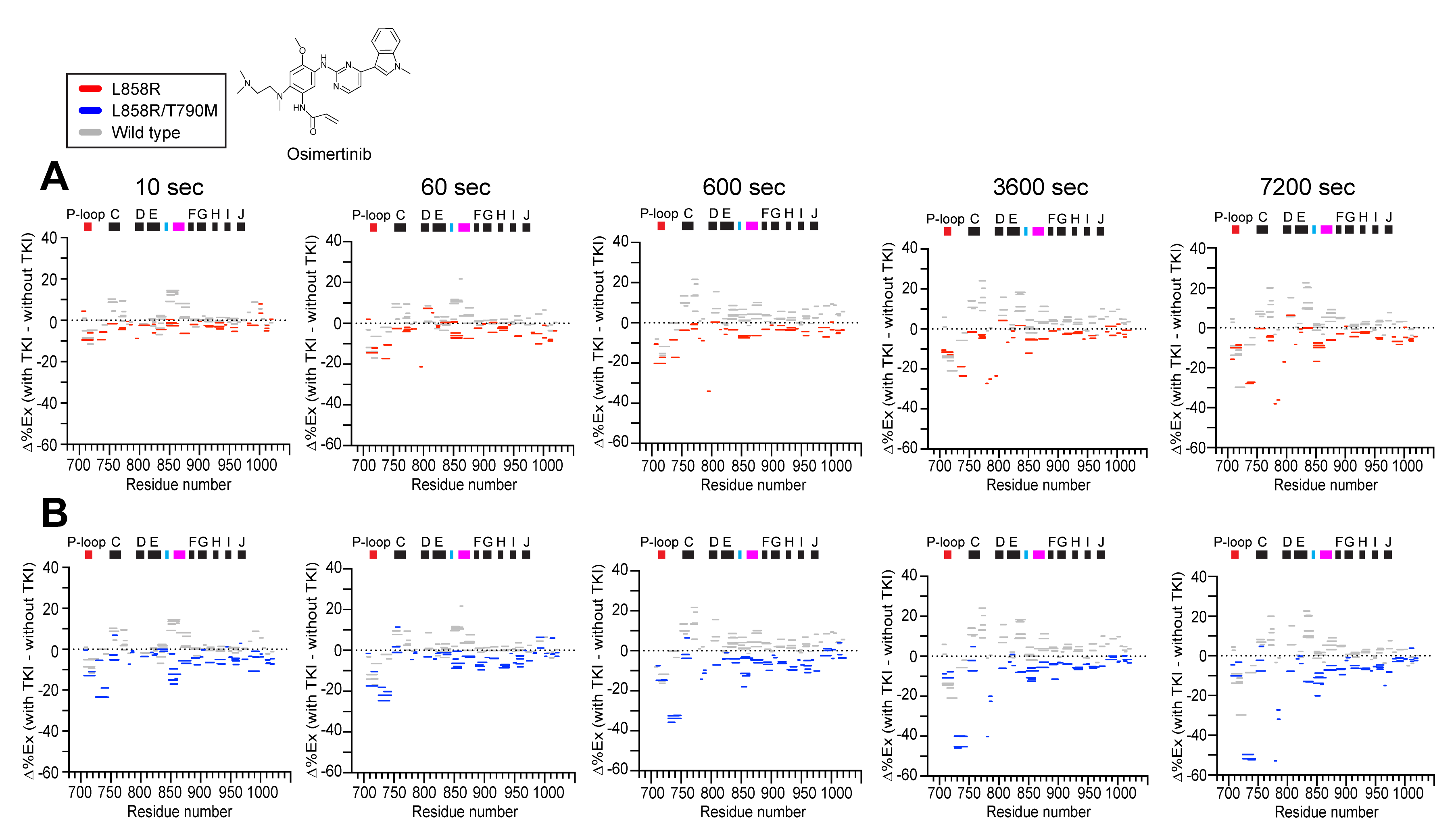

**Figure S22: Woods Plots of osimertinib-EGFR L858R complexes**

Woods plots (Δ%EX map) for the EGFR TKD complexed with covalent osimertinib at the deuterium labeling time of 10, 60, 600, 3600, and 7200 sec at 25^o^C, pD 7.4. The percentage exchange differences between the TKI-bound and -unbound TKD (Δ%Ex) of all analyzed peptides, shown as colored horizontal bars, are plotted with the residue number on the X-axis. The regions with positive or negative Δ%Ex values become structurally more flexible or rigid, respectively, with indicated TKIs relative to the TKI-unbound TKD. The position of α-helices (black bar) and functionally important regions, including P-loop, HRD (cyan bar), and DFG·A-loop (magenta bar), are shown at the top of each Δ%EX map. All HDX data were obtained from n = 3 separate protein preparations, with three independent experiments for each. HDX data statistics are presented in Tables S6 and S7.

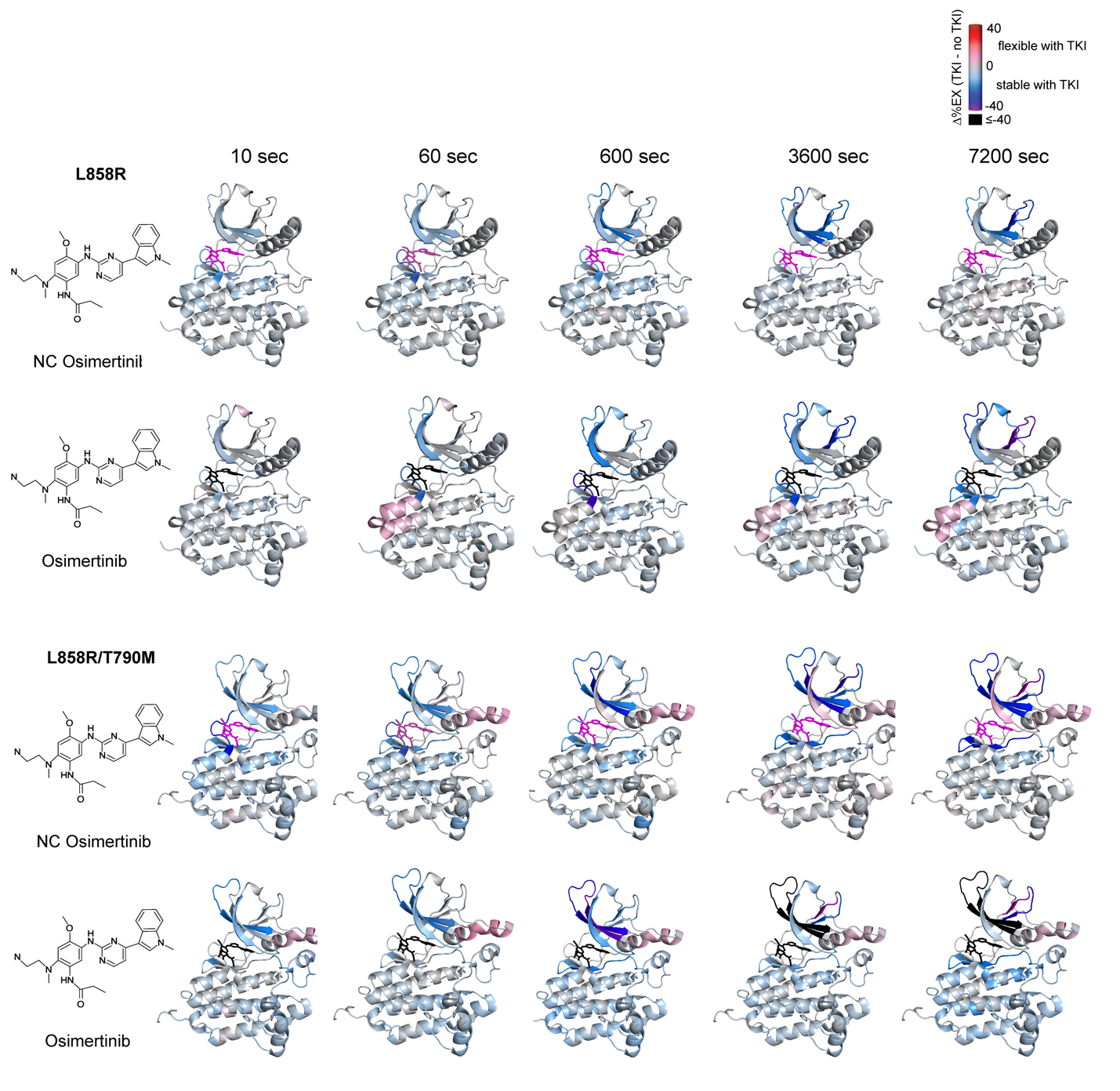

**Figure S23: Structural dynamics of the EGFR L858R or L858R/T790K mutant complexed with non-covalent osimertinib or osimertinib**

The Δ%Ex values are color-coded and mapped onto crystal structures of L858R (PDB ID: 6JWL)^48^ and L858R/T790M (PDB ID: 6JXO).^48^ The bound inhibitor is indicated as a stick in each figure. All HDX data were obtained from n = 3 separate protein preparations, with three independent experiments for each. The figures are generated using PyMOL (http://www.pymol.org).

**Table S1. The HDX data statistics for the wild type EGFR TKD without and with the first-generation TKIs.**

| **Data Set** | No TKI | Erlotinib | Gefitinib | Lapatinib |
| --- | --- | --- | --- | --- |
| HDX reaction details | 20 mM HEPES, 100 mM NaCl (pD 7.4) at 25^o^C | | | |
| HDX time course (sec) | 0, 10, 60, 600, 3600, 7200 | | | |
| HDX control samples | 20 mM HEPES, 100 mM NaCl, 8 M Urea-d4 (pD 7.4) at 25^o^C | | | |
| Back-exchange (%) | 47.4 | | | |
| # of Peptides | 138 | 120 | 83 | 100 |
| Sequence coverage (%) | 97.2 | 94.4 | 94.1 | 87.9 |
| Average peptide length / Redundancy | 10.7/4.4 | 10.1/4.35 | 10.6/3.74 | 10.1/3.63 |
| Replicates (biological and technical) | 3 technical repeats on each of 3 separate protein preparations | | | |
| Repeatability (Ave Standard deviation)* | 2.4 % (0.12 Da) | 1.8 % (0.09 Da) | 2.5 % (0.13 Da) | 2.5 % (0.11 Da) |
| Significant differences in HDX  (TKI- no TKI)** | N/A | 4.9 % (0.24 Da) | 7.7 % (0.39 Da) | 7.9 % (0.28 Da) |

*Average standard deviation (SD) calculated based on the deuterium uptake of biological repeat experiments (3 separate protein preparations, n=3).

**Significance in % (and Da) at 98 % confidence calculated based on Houde *et al*. (2011) J of Pharm Sci 100, 2071-2086.

**Table S2. The HDX data statistics for the wild type EGFR TKD with the second-generation TKIs.**

| **Data Set** | NC Afatinib | NC Dacomitinib | NC Osimertinib |
| --- | --- | --- | --- |
| HDX reaction details | 20 mM HEPES, 100 mM NaCl (pD 7.4) at 25^o^C | | |
| HDX time course (sec) | 0, 10, 60, 600, 3600, 7200 | | |
| HDX control samples | 20 mM HEPES, 100 mM NaCl, 8 M Urea-d4 (pD 7.4) at 25^o^C | | |
| Back-exchange (%) | 47.4 | | |
| # of Peptides | 77 | 72 | 94 |
| Sequence coverage (%) | 94.4 | 91.7 | 97.3 |
| Average peptide length / Redundancy | 10.8/3.88 | 10.6/3.38 | 11.3/4.93 |
| Replicates (biological and technical)* | n = 3 | n = 2 | n=3 |
| Repeatability (Ave Standard deviation)** | 3.3 % (0.16 Da) | 2.2 % (0.11 Da) | 2.1 (0.11 Da) |
| Significant differences in HDX  (TKI- no TKI)*** | 9.2 % (0.46 Da) | 7.1 % (0.34 Da)^#^ | 7.0 % (0.35 Da) |

*3 technical repeats on each of 3 separate protein preparations (n=3) except for NC dacomitinib experiments (3 technical repeats on each of 2 separate protein preparations, n=2)

** Average standard deviation (SD) calculated based on the deuterium uptake of biological repeat (n = 2 or 3) experiments

***Significance in % (or Da) at 98 % confidence calculated based on Houde *et al*. (2011) J of Pharm Sci 100, 2071-2086.

\# Significance in % (or Da) at 92 % confidence

**Table S3. The HDX data statistics for the wild-type EGFR TKD with compound 2 or 3.**

| **Data Set** | Compound 2 | Compound 3 |
| --- | --- | --- |
| HDX reaction details | 20 mM HEPES, 100 mM NaCl (pD 7.4) at 25^o^C | |
| HDX time course (sec) | 0, 10, 60, 600, 3600, 7200 | |
| HDX control samples | 20 mM HEPES, 100 mM NaCl, 8 M Urea-d4 (pD 7.4) at 25^o^C | |
| Back-exchange (%) | 47.4 | |
| # of Peptides | 77 | 106 |
| Sequence coverage (%) | 96.7 | 97.6 |
| Average peptide length / Redundancy | 11.0/2.81 | 10.9/3.94 |
| Replicates (biological and technical)* | 3 | 2 |
| Repeatability (Ave Standard deviation)** | 2.1 % (0.11 Da) | 2.4 % (0.12 Da) |
| Significant differences in HDX  (TKI- no TKI)*** | 7.1 % (0.36 Da) | 9.9 % (0.48 Da)^#^ |

*3 technical repeats on each of 3 separate protein preparations (n=3) for compound 2 experiments, n=2 for compound 3 experiments (3 technical repeats on each of 2 separate protein preparations, n=2)

** Average standard deviation (SD) calculated based on the deuterium uptake of biological repeat (n = 2 or 3) experiments

***Significance in % (or Da) at 98 % confidence calculated based on Houde *et al*. (2011) J of Pharm Sci 100, 2071-2086.

\# Significance in % (or Da) at 90 % confidence

**Table S4: Crystallization Conditions, Data Collection and Refinement Statistics**

| Protein(s) | EGFR-TKD | EGFR-TKD |
| --- | --- | --- |
| Residue boundaries | 696-1022 | 696-1022 |
| Co-crystallized or soaked | Co-crystallized | Soaked |
| inhibitor | Non-covalent osimertinib | Non-covalent osimertinib |
| PBD accession code | ---- | ---- |
| Crystallization Conditions | protein (5 mg/mL, 20 mM Tris, pH 7.5, 250 mM NaCl, 2.5% 1,3-propanediol) : reservoir solution (100 mM MES, pH 6.0, 0.8 M sodium citrate) ratio = 1:1, 16 ºC | protein (5.5 mg/mL, 20 mM Tris, pH 7.5, 250 mM NaCl, 2.5% 1,3-propanediol) : reservoir solution (100 mM HEPES, pH 7.0, 0.6-0.9 Na-K tartrate) ratio = 1:1, 16 ºC |
| Data Collection^a^ | | |
| Beamline | APS/NE-CAT 24-ID-E | APS/NE-CAT 24-ID-E |
| Date of collection | October 15, 2019 | February 13, 2019 |
| Wavelength (Å) | 0.97918 | 0.97918 |
| Space Group | *I 2 3* | *I 2 3* |
| *Cell Dimensions* | | |
| a, b, c (Å) | 145.97, 145.97, 145.97 | 145.92, 145.92, 145.92 |
| α, β, γ (Å) | 90.00, 90.00, 90.00 | 90.00, 90.00, 90.00 |
| Resolution (Å) | 103.2 – 2.3 | 51.59 – 2.55 |
| Completeness | 99.9 (98.8) | 100.0 (99.9) |
| R_sym_ | 0.152 (2.1) | 0.075 (6.05) |
| I/σ | 18.5 (1.3) | 16.0 (2.7) |
| CC^1/2 b^ | 0.986 (0.81) | 0.996 (0.32) |
| Refinement | | |
| Number of reflections | 22,884 (2794) | 16,995 (2807) |
| R_work_/R_free_ (%) | 20.2/22.5 | 20.6/24.5 |
| *Number of atoms* | | |
| Protein | 2527 | 2492 |
| Ions | 1 | 0 |
| Ligands | 51 | 42 |
| Water | 64 | 2 |
| *Average B factor (Å)* | | |
| Protein | 88.6 | 96.14 |
| Ions | - | - |
| Ligands | 95.55 | 104.36 |
| Water | 82.53 | 71.03 |
| *Geometry (Ramachandran)* | | |
| Favored (%) | 96.8 | 96.1 |
| Allowed (%) | 2.9 | 3.6 |
| Outliers (%) | 0.3 | 0.3 |
| *RMSD (Å)* | | |
| Bond length | 0.005 | 0.002 |
| Bond angle | 0.8 | 0.62 |

^a^Numbers in parentheses denote the highest resolution shell

^b^CC^1/2^ reported for the highest resolution shell

**Table S5. The HDX data statistics for wild-type EGFR TKD covalent complexes**

| **Data Set** | Afatinib | Dacomitinib | Osimertinib |
| --- | --- | --- | --- |
| HDX reaction details | 20 mM HEPES, 100 mM NaCl (pD 7.4) at 25^o^C | | |
| HDX time course (sec) | 0, 10, 60, 600, 3600, 7200 | | |
| HDX control samples | 20 mM HEPES, 100 mM NaCl, 8 M Urea-d4 (pD 7.4) at 25^o^C | | |
| Back-exchange (%) | 47.4 | | |
| # of Peptides | 71 | 78 | 79 |
| Sequence coverage (%) | 92.6 | 93.2 | 91.4 |
| Average peptide length / Redundancy | 10.4/3.19 | 9.5/3.23 | 11.5/3.71 |
| Replicates (biological and technical)* | n = 3 | n = 2 | n=3 |
| Repeatability (Ave Standard deviation)** | 2.4 % (0.10 Da) | 1.7 % (0.079 Da) | 1.4 % (0.070 Da) |
| Significant differences in HDX  (TKI- no TKI)*** | 7.5 % (0.34 Da) | 10.5 % (0.47 Da)^#^ | 5.9 % (0.30 Da) |

*3 technical repeats on each of 3 separate protein preparations (n=3) except for dacomitinib experiments (3 technical repeats on each of 2 separate protein preparations, n=2)

**Average standard deviation (SD) calculated based on the deuterium uptake of biological repeat (n = 2 or 3) experiments ***Significance in % (or Da) at 98 % confidence calculated based on Houde *et al*. (2011) J of Pharm Sci 100, 2071-2086.

\# Significance in % (or Da) at 90 % confidence

**Table S6. The HDX data statistics for the L858R TKD encounter and covalent complexes**

| **Data Set** | L858R (No TKI) | L858R (NC Osimertinib) | L858R (Osimertinib) |
| --- | --- | --- | --- |
| HDX reaction details | 20 mM HEPES, 100 mM NaCl (pD 7.4) at 25^o^C | | |
| HDX time course (sec) | 0, 10, 60, 600, 3600, 7200 | | |
| HDX control samples | 20 mM HEPES, 100 mM NaCl, 8 M Urea-d4 (pD 7.4) at 25^o^C | | |
| Back-exchange (%) | 37.7 % | | |
| # of Peptides | 67 | 64 | 61 |
| Sequence coverage (%) | 92.6 | 88.5 | 87.3 |
| Average peptide length / Redundancy | 10.7/2.61 | 10.8/2.37 | 12.4/2.59 |
| Replicates (biological and technical) | 3 technical repeats on each of 3 separate protein preparations (n=3) | | |
| Repeatability (Ave Standard deviation)* | 1.7 % (0.11 Da) | 2.0 (0.10 Da) | 2.3 % (0.14 Da) |
| Significant differences in HDX  (TKI- no TKI)** | N/A | 5.9 % (0.32 Da) | 6.6 % (0.39 Da) |

*Average standard deviation (SD) calculated based on the SD of biological repeat experiments (3 separate protein preparations, n=3).

**Significance in % (or Da) at 98 % confidence calculated based on Houde *et al*. (2011) J of Pharm Sci 100, 2071-2086.

**Table S7. The HDX data statistics for the L858R/T790M TKD encounter and covalent complexes**

| **Data Set** | L858R/T790M  (No TKI) | L858R/T790M (NC Osimertinib) | L858R/T790M (Osimertinib) |
| --- | --- | --- | --- |
| HDX reaction details | 20 mM HEPES, 100 mM NaCl (pD 7.4) at 25^o^C | | |
| HDX time course (sec) | 0, 10, 60, 600, 3600, 7200 | | |
| HDX control samples | 20 mM HEPES, 100 mM NaCl, 8 M Urea-d4 (pD 7.4) at 25^o^C | | |
| Back-exchange (%) | 40.7 | | |
| # of Peptides | 67 | 78 | 92 |
| Sequence coverage (%) | 91.7 | 90.8 | 90.8 |
| Average peptide length / Redundancy | 11.8/2.55 | 11.6/3.14 | 12.2/3.75 |
| Replicates (biological and technical) | 3 technical repeats on each of 3 separate protein preparations (n=3) | | |
| Repeatability (Ave Standard deviation)* | 2.8 % (0.16 Da) | 2.1 % (0.12 Da) | 2.3 % (0.14 Da) |
| Significant differences in HDX  (TKI- no TKI)** | N/A | 7.5 % (0.44 Da) | 7.8 % (0.47 Da) |

*Average standard deviation (SD) calculated based on the deuterium uptake of biological repeat experiments (3 separate protein preparations, n=3).

**Significance in % (or Da) at 98 % confidence calculated based on Houde *et al*. (2011) J of Pharm Sci 100, 2071-2086.
